# Structural and molecular basis of FAN1 defects in promoting Huntington’s disease

**DOI:** 10.1101/2024.10.07.617005

**Authors:** F. Li, A. Phadte, M. Bhatia, S. Barndt, A. R. Monte Carlo, C-F. D. Hou, R. Yang, S. Strock, A. Pluciennik

## Abstract

FAN1 is a DNA dependent nuclease whose proper function is essential for maintaining human health. For example, a genetic variant in FAN1, Arg507 to His hastens onset of Huntington’s disease, a repeat expansion disorder for which there is no cure. How the Arg507His mutation affects FAN1 structure and enzymatic function is unknown. Using cryo-EM and biochemistry, we have discovered that FAN1 arginine 507 is critical for its interaction with PCNA, and mutation of Arg507 to His attenuates assembly of the FAN1-PCNA on a disease-relevant extrahelical DNA extrusions formed within DNA repeats. This mutation concomitantly abolishes PCNA-FAN1-dependent cleavage of such extrusions, underscoring the importance of PCNA to the genome stabilizing function of FAN1. These results unravel the molecular basis for a specific mutation in FAN1 that dramatically hastens the onset of Huntington’s disease.

## INTRODUCTION

DNA repair systems have evolved to rectify exogenous or endogenous DNA lesions. FAN1 (FANCD2/FANCI associated nuclease 1)^1-5^ is a DNA repair enzyme involved in removal of extrahelical extrusions formed within triplet repeat sequences, and DNA interstrand crosslinks (ICL) at stalled replication forks^1,4-8^. Genetic variants in the human FAN1 gene are associated with modification of age of onset of Huntington’s disease (HD)^9-11^, a neurodegenerative disorder caused by expansion of a CAG repeat tract within exon 1 of the huntingtin gene^12^. In model systems, knockout/knockdown of FAN1 promotes triplet repeat expansion ^10,11,13-17^. Because expansion of CAG repeats in striatal and cortical neurons over the lifetime of an individual is a key driver of HD manifestation^18^, FAN1 is regarded as playing a preventative role in HD by maintaining the genetic stability of triplet repeats^19^. Moreover, FAN1 has also been implicated in stabilization of triplet repeats associated with fragile X related disorders^13,17^, suggesting that FAN1 may have a broader neuroprotective role from repeat expansions in humans.

FAN1 is a 1017-amino-acid protein that harbors a RAD18-like ubiquitin-binding zinc finger (UBZ) domain near the N-terminus, an SAF-A/B, Acinus, and PIAS (SAP)-type primary DNA binding domain (in the central region), a tetratricopeptide repeat (TPR) domain, and a virus-type replication-repair nuclease module (VRR_nuc) domain in the C-terminal region^19,20^. The FAN1 enzyme possesses 5’flap endonuclease and 5’ to 3’ exonuclease activities ^1,2,4,5^ (mediated by its VRR_nuc domain) and was originally identified as a key player in the reversal of interstrand crosslink (ICL) damage. Indeed, inactivation of FAN1 results in increased sensitivity of human cells to treatment with ICL-inducing agents like mitomycin C or cisplatin ^1,3,4^, and in humans leads to karyomegalic interstitial nephritis- a chronic kidney disease^21^, and autism/schizophrenia^22^. We have recently shown that the endonuclease and exonuclease functions of FAN1 act in concert to cleave extrahelical extrusions (structures that arise due to DNA strand slippage within repetitive sequences) by a process that requires DNA-loaded sliding clamp proliferating cell nuclear antigen (PCNA)^6^. The FAN1-catalyzed cleavage of such structures leads to their removal by factors that are yet to be identified. The activation of the FAN1 nuclease function by PCNA is a strand-directed process wherein the orientation of DNA-loaded PCNA has been suggested to orient the nuclease activity of FAN1 to a particular DNA strand. This mechanism requires a physical interaction between these proteins on DNA, although the molecular events underlying this activation process are not understood.

In this work, we have determined the cryo-EM structure of the FAN1-PCNA complex clamped to a CAG extrahelical extrusion. Using a combination of structural and biochemical approaches, we show that the HD onset-modifying R507H variant of FAN1 abolishes its PCNA-dependent stimulation as a result of structural distortions in the FAN1-PCNA binding interface. These data reveal that disruption of PCNA-dependent stimulation of FAN1 via R507 mutation is responsible for promoting the rapid onset of HD.

## RESULTS

### Cryo-EM structure of FAN1-PCNA-DNA ternary complex

FAN1 is a DNA repair nuclease that can act on a wide range of DNA substrates. We have recently shown that FAN1 nuclease activity is triggered by small extrahelical extrusions (structures that arise from DNA strand slippage within repetitive sequences during helix opening)^6^. FAN1 nuclease activity is stimulated by the DNA-loaded sliding clamp PCNA to cleave DNA substrates harboring triplet repeat extrahelical extrusions at physiological ionic strength^6^ (Fig. 1a-c and Extended Data Fig. 1a-c). Fine mapping of PCNA-activated FAN1 nuclease cleavage of a linear DNA substrate harboring a (CAG)_2_ extrahelical extrusion indicates that the predominant endonucleolytic cleavage occurs on the strand harboring the DNA extrusion 2 nucleotides 3’ to the extrusion (resulting in 18 nucleotides cleavage product), followed by nucleolytic cuts giving rise to products of 15, 13, 10, 9, and 8 nucleotides becoming evident as a function of time (Fig. 1a-c and Extended Data Fig. 1b,c). This periodicity is reminiscent of previous observations of patterns of FAN1 cleavage of flap-containing DNAs^7^. To further understand the function of PCNA in this reaction, we used cryogenic-electron microscopy (cryo-EM) to examine the assembled FAN1-PCNA-DNA ternary complex, which contains recombinant full-length human proteins FAN1 and PCNA and a linear DNA substrate harboring a (CAG)_2_ extrahelical extrusion (Extended Data Table 1). The complex was isolated by size exclusion chromatography and vitrified on grids for cryo-EM data collection (Extended Data Fig. 2). Three major classes of the FAN1-PCNA-DNA complex were identified using single particle analysis (SPA) (Extended Data Fig. 3-5). The most complete of these was class 1, which yielded a 3D map at 3.8 Å resolution at a 0.143 cut-off Fourier Shell Correlation (FSC) (Extended Data Fig. 5a,d). The map was further sharpened using DeepEMhancer^23^ (Fig. 1d). We used AlphaFold 3^24^ predicted models of FAN1, PCNA and DNA as the initial templates for model building according to the sharpened maps. After real-space refinement, the models achieved a final correlation coefficient (CC) of approximately 0.82, indicating an excellent fit between the model and the density map (Table 1). The other two 3D class maps are resolved at 3.4 Å and 5.9 Å for class 2 and 3, respectively (Extended Data Fig. 5b,c). The local resolution estimation of these three classes was also performed (Extended Data Fig. 5d-f).

**Figure 1.**
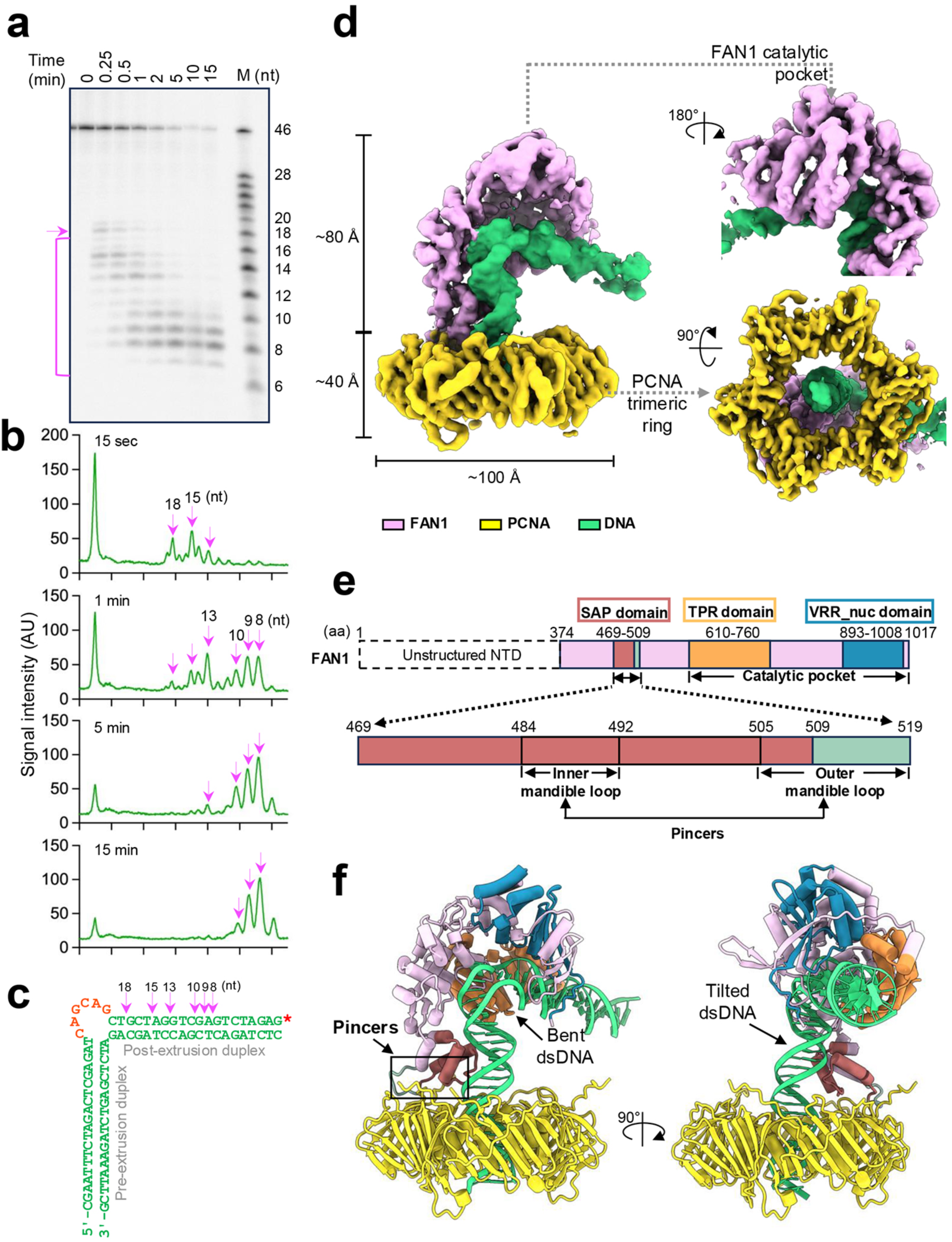
PCNA-dependent FAN1 nuclease activity and the cryo-EM structure of FAN1-PCNA-DNA complex. (**a**) A time course of PCNA-dependent FAN1 nuclease activity on a 3’-radiolabeled linear DNA substrate harboring a (CAG)_2_ extrahelical extrusion. FAN1 endonucleolytic cleavage is marked by the pink arrow, and cleavage products are indicated by the pink bracket. The 5’ radiolabeled DNA marker has the same nucleotide composition as the DNA strand harboring the extrusion. (**b**) Densitometric scans of the gel above at the indicated time points show the sequential appearance of cleavage products (pink arrows). (**c**) Schematic of the DNA substrate used in this experiment, with the major FAN1 cleavage sites depicted by pink arrows. The red asterisk indicates the 3’-radiolabeled DNA end. The numbers represent the length of the products post FAN1 cleavage. (**d**) FAN1-PCNA-DNA ternary complex cryo-EM maps showing the complete complex from a front view (left), FAN1 catalytic pocket from a back view (upper right), and PCNA trimeric ring from a bottom view (bottom right). (**e**) Schematic representation of the human FAN1 protein (as shown previously^19^) with domains demarcated in different colors. (**f**) Atomic model of the FAN1-PCNA-DNA ternary complex, with FAN1 domains colored as per the schematic in panel (**e**). See also Extended Data Fig. 1-5 and Table 1.

**Table 1.**
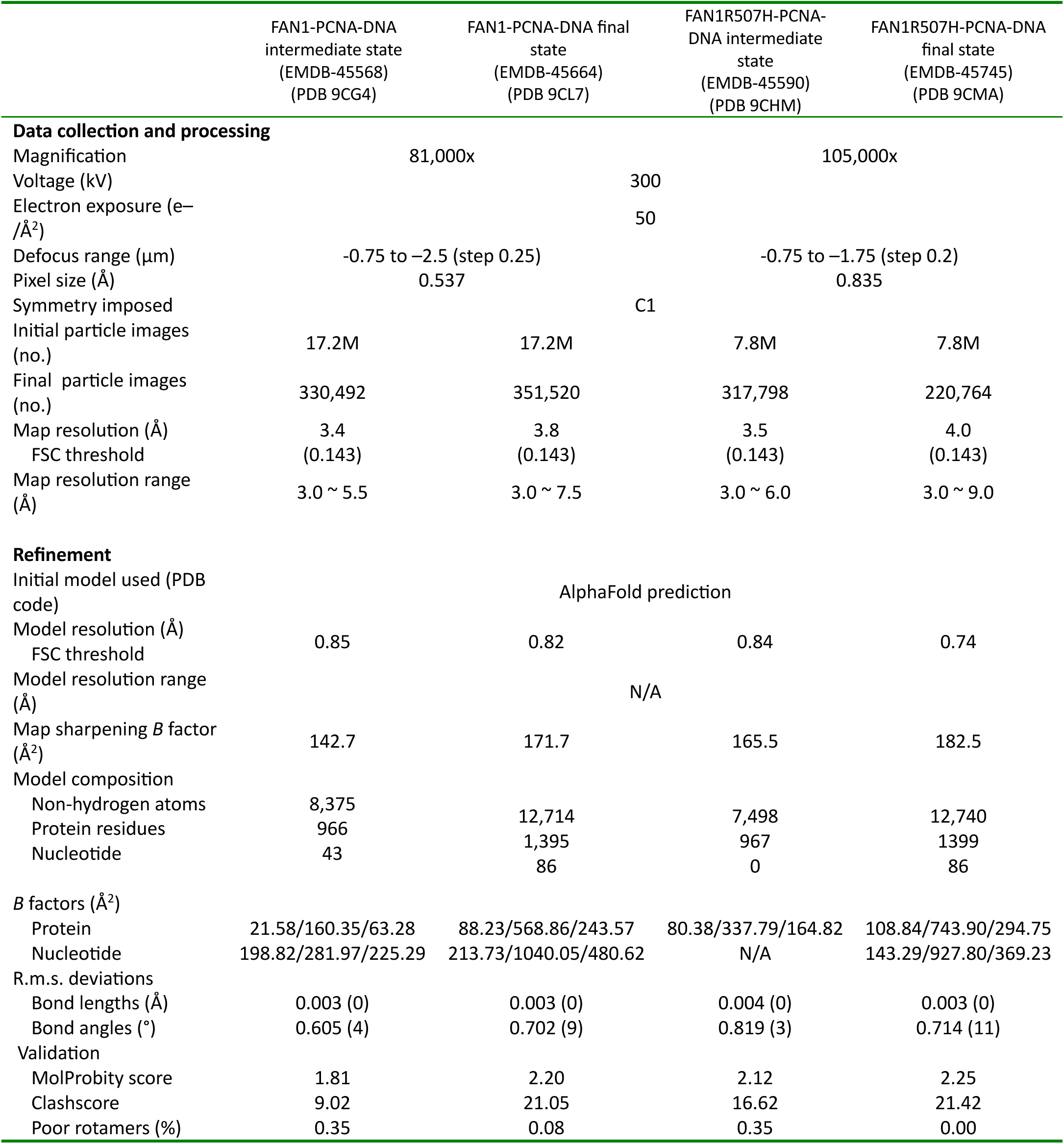
Cryo-EM data collection, refinement and validation statistics.

The FAN1-PCNA-DNA complex (3D class 1) has approximate dimensions of 120 Å x 100 Å x 100 Å and reveals a FAN1 monomer bound to a PCNA trimeric ring that encircles the double-stranded DNA substrate harboring a (CAG)_2_ extrusion. The DNA substrate is threaded through the PCNA ring at a tilted angle relative to the vertical axis of trimeric ring surface (Fig. 1d,f). FAN1 also makes contact with the DNA substrate using its catalytic pocket (including TPR and VRR-nuclease domains), and the DNA is bent at the extrahelical extrusion (Fig. 1d,e). FAN1 interacts with one of the three PCNA protomers and extends perpendicularly from the p21-binding face of the PCNA ring^25^ (Fig. 1f). It is noteworthy in this regard that the N-terminal region of FAN1 (1-373) is not visible in our cryo-EM maps even though the full length human FAN1 was used, suggesting that the N-terminal region is flexible, in agreement with previous reports^7^. The cryo-EM map of class 2 is highly similar to the class 1 map, showing the same spatial arrangement of the three components, but some regions are more anisotropic. This class exhibits a less resolved catalytic pocket of FAN1, suggesting reduced stability of the interaction between the catalytic domain and the DNA extrusion (Extended Data Fig. 5e). The class 3 map shows a helical DNA protruding from the bottom of the PCNA ring such that the protein is positioned closer to the vertical midpoint of the DNA helical axis. The other end of the DNA is enveloped by density that can accommodate two α-helices of the FAN1 SAP domain, suggestive of DNA binding by FAN1 (Extended Data Fig. 5f). The three classes observed here may represent intermediate states of the FAN1-PCNA-DNA complex during substrate recognition and cleavage (see Discussion).

### FAN1 clips onto two back-to-back hydrophobic pockets on PCNA

The interaction interface between FAN1 and PCNA as evident in both class 1 and 2 maps is defined by “pincers” in FAN1 serving as the primary binding site with one PCNA protomer (Fig. 1e,f). This pincer architecture is comprised of two flexible “mandibles”: an inner mandible loop corresponding to amino acids 484-492 within the SAP domain, and an outer mandible loop comprising of amino acids 505-519, which partially overlaps with the SAP domain (Fig.1e and Fig. 2a,b). Coulombic surface potential analyses revealed a positively charged area (pI: 10.53) on the pincers of FAN1 and a highly negatively charged area (pI: 3.89) on the outer side of the PCNA ring, which mediates electrostatic attraction and thus enables the interaction between FAN1 and PCNA (Supplemental Data Fig. 1). The two mandible loops of FAN1 interact with two distinct back-to-back hydrophobic pockets of PCNA (Fig. 2c), with the interface area measuring approximately 775:816 Å². The inner hydrophobic pocket of PCNA, which is formed by amino acids L22, Y211, and F214, establishes hydrophobic contacts with the FAN1 inner mandible loop (Fig. 2c,d). Likewise, the outer hydrophobic pocket of PCNA, consisting of amino acids M40, V45, L47, I126, I128, V233, L251, A252, I255, Y133, and Y250, engages with the FAN1 outer mandible loop residues (Fig. 2c,e).

**Figure 2.**
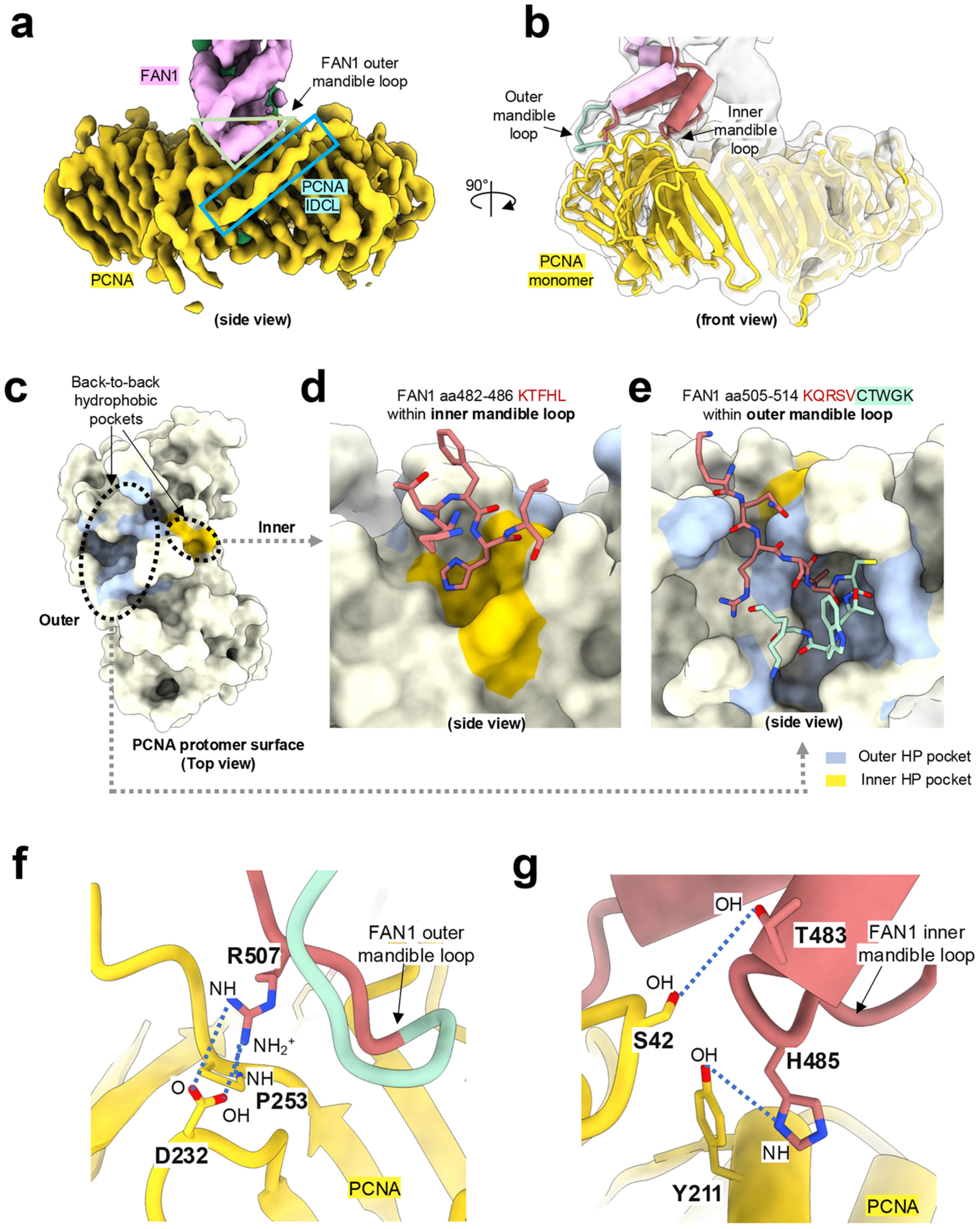
The interaction between FAN1 and PCNA. (**a**) Cryo-EM map showing a side view of FAN1 and PCNA interaction sites. (**b**) Atomic model (front view overlaid with a density map) of FAN1 inner and outer mandible loops that clip one PCNA monomer. (**c**) PCNA protomer surface (top view), showing two back-to-back hydrophobic (HP) pockets. (**d**) Interaction of FAN1 aa 482-486 within the inner mandible loop with the inner hydrophobic pocket of PCNA composed of residues: (1) Aliphatic: L22. (2) Aromatic: Y211 and F214. (**e**) Interaction of FAN1 aa 505-514 within outer mandible loop with the outer hydrophobic pocket of PCNA consisting of residues: (1) Aliphatic: M40, V45, L47, I126, I128, V233, L251, A252 and I255. (2) Aromatic Y133 and Y250. (**f**) Hydrogen bonds between PCNA and the FAN1 outer mandible loop. (**g**) Hydrogen bonds between the FAN1 inner mandible loop and PCNA. See also Extended Data Fig. 6 and Extended Data Table 2.

Further analysis of the FAN1-PCNA interaction interface using the PDBsum^26^ program identified multiple interactions contributing to the stability of this complex, including four hydrogen bonds, one salt bridge, and 74 van der Waals interactions (Extended Data Table 2). Notably, Arg507 on FAN1 forms a salt bridge and a hydrogen bond with residue Asp232, as well as an additional hydrogen bond with Pro253 on PCNA (Fig. 2f). Furthermore, the His485 and Thr483 on FAN1 establish hydrogen bonds with Tyr211 and Ser42 on PCNA, respectively (Fig. 2g). These intricate interactions underscore the critical role of the pincers in mediating the stable association between FAN1 and PCNA, which is essential for the activation of FAN1 extrusion cleavage activity by PCNA.

### Interaction interface of PCNA with DNA

Prior structural, single-molecule, and molecular dynamics studies have suggested that movement of the sliding clamp on DNA is facilitated by a continuous tilt and rotate mechanism, with the tilt angles of DNA ranging from 15° to 40°^27-30^. Both of our cryo-EM structures (class 1 and 2) reveal a DNA molecule tilted at ∼15° from the Y axis in the Y-Z plane (Extended Data Fig. 6a,c). PCNA makes several contacts with DNA, although the 3 subunits do not contribute equally to the PCNA-DNA interaction interface (Fig. 3). Additionally, based on the comparison of the class 1 and 2 complexes, we observe an angular displacement of the DNA by ∼4° in the X-Y plane when the complex is in its most stable conformation (class 1) (Extended Data Fig. 6b). These states may be reflective of movement of the PCNA clamp on the DNA via a cog-wheel mechanism as has been previously proposed^27^ (see Discussion).

**Figure 3.**
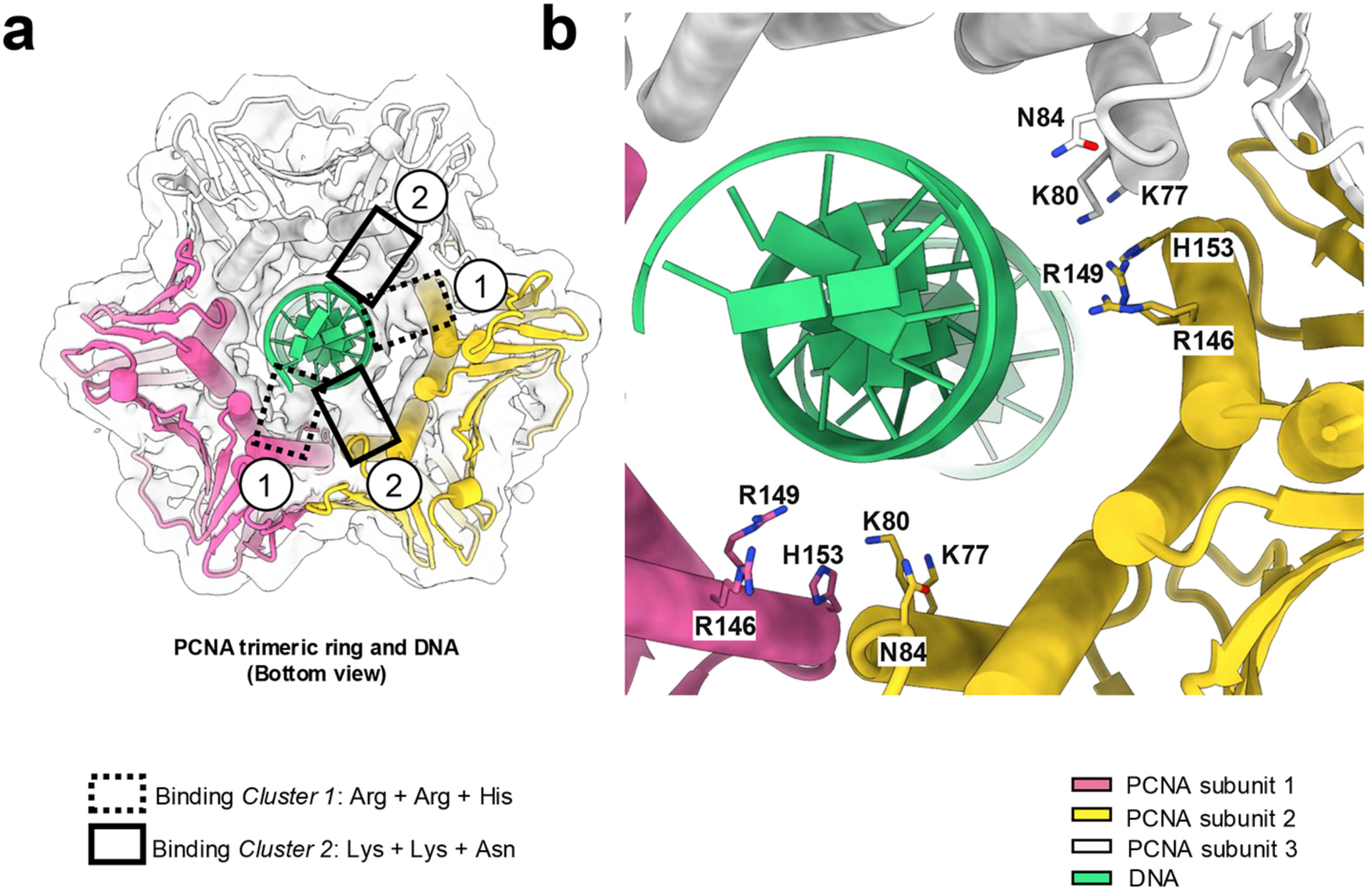
The PCNA-DNA interface. (**a**) Density map showing four interaction clusters between the PCNA trimeric ring and dsDNA. These clusters are asymmetric with respect to the DNA axis and are biased to one side of the PCNA ring. The PCNA subunits shown in pink and white contact DNA via *Cluster 1* and *Cluster 2* residues, respectively, whereas the PCNA subunit indicated in yellow makes contacts via both *Cluster 1* and *Cluster 2* residues. (**b**) *Cluster 1* residues: Arg146 + Arg149 + His153; *Cluster 2* residues: Asn84 + Lys80 + Lys77. See also Extended Data Fig. 6.

### Role of FAN1 major domains in interaction with extrahelical extrusion-containing DNA

The electrostatic surface potential of FAN1 reveals a positively charged surface that extends from the SAP domain to the catalytic pocket, facilitating DNA binding (Supplemental Data Fig. 1). In our cryo-EM analysis, the class 1 map reveals density corresponding to the entirety of the DNA substrate, with the DNA bent at the catalytic pocket of FAN1 by an inner angle of 78° (Fig. 1f). The bent conformation of the DNA is consistent with previous studies of extrusion-containing DNAs^31-34^, and is reminiscent of the bent double flap DNA substrate described previously in a co-crystal structure with a truncated FAN1 variant (PDB: 4RI8)^7^. It should be noted here that DNAs harboring extrahelical extrusions are naturally bent^31-33^, a conformational feature that may facilitate recognition by FAN1. The FAN1 SAP domain contacts the pre-extrusion DNA duplex via amino acids S473, K493, and K482, stabilizing the DNA in the vicinity of the PCNA ring (Fig. 4a,c). The region between the SAP and TPR domains within the catalytic pocket of FAN1 accommodates the (CAG)₂ extrusion, and involves amino acids Y374, R420, Y436, R424, and K425, providing an extensive recognition interface for extrusion sensing (Fig. 4a,b). The helical repeats of the TPR domain collaborate with the insertion loop of the VRR_nuc domain to grip the DNA, using amino acids R679, H681, H716, H718, R752, and R982, securing the DNA at a fixed angle beyond the extrusion site (Fig. 4d-f).

**Figure 4.**
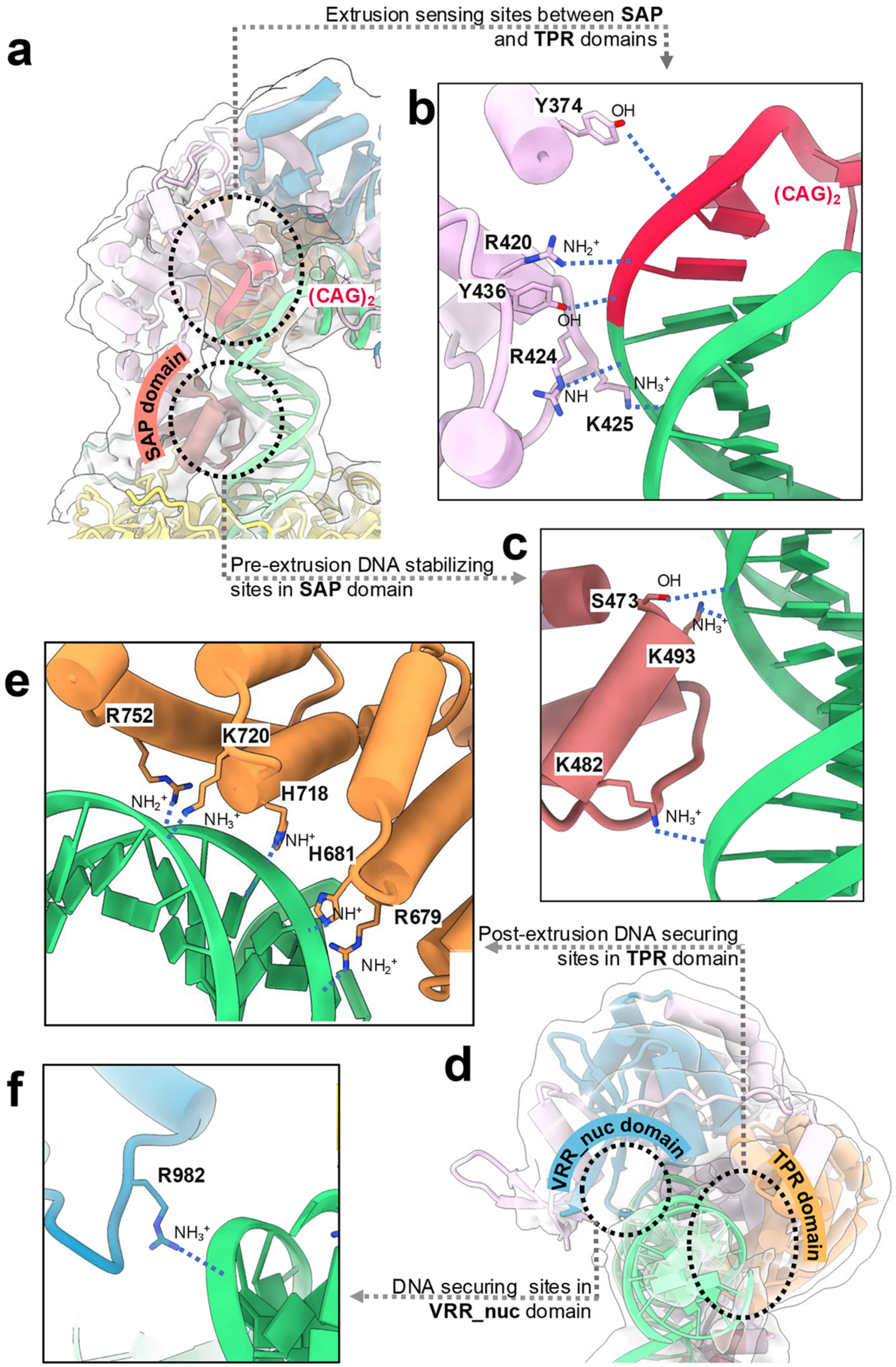
Interaction between FAN1 and DNA. (**a**) Density map of the FAN1-DNA interface overlaid with atomic model. (**b**) Residues S473, K493, and K482 from the region connecting the SAP and TPR domains function as extrusion sensing sites. (**c**) FAN1 SAP domain stabilizes the DNA via residues Y374, R420, R424, K425, and Y436. (**d**) DNA helical axial view of density map of the FAN1-DNA interface overlaid with atomic model. (**e-f**) DNA securing contacts made by residues in the TPR domain (R679, H681, H718, K720, and R752), and the VRR_nuc domain (R982).

We have shown previously that the PCNA-activated FAN1 nuclease displays exquisite specificity for extrusion-containing DNAs, with little to no activity observed on homoduplex DNA^6^. To establish the structural basis, if any, of this feature of FAN1 behavior, we determined the cryo-EM structure of FAN1-PCNA complex bound to a homoduplex DNA (Extended Data Fig. 7). We observe two distinct properties of the homoduplex-containing ternary complex: (i) The DNA homoduplex is linear and displays no bends, and (ii) FAN1 is bound to the DNA end, effectively capping it. We postulate that the lack of a DNA bend does not allow FAN1 to capture the DNA substrate in a conformation conducive to catalysis.

### R507 in FAN1 is critical for PCNA-stimulated FAN1 nuclease activity

Genome wide association studies of HD patients have identified single nucleotide polymorphisms in the *FAN1* gene as modifiers of disease onset, with the R507H variant associated with hastening HD onset by 5.2 years^9^. Attempts to understand the functional consequence of the R507H mutation using cellular approaches have been inconclusive^10,11^. As noted above, our cryo-EM data revealed a key role played by Arg507 on FAN1 in mediating the FAN1-PCNA interaction. This residue interacts with Asp232 on PCNA via a salt bridge and a hydrogen bond, and also with Pro253 on PCNA through a hydrogen bond (Fig. 2f,g). Therefore, we hypothesized that mutation of FAN1 Arg507 might impact the PCNA-FAN1 interaction, and thereby affect the ability of PCNA to activate FAN1. Hence, we mutated and purified recombinant FAN1 harboring an Arg507 to His substitution; likewise, we also prepared recombinant FAN1 with an Arg507 to Ala substitution to completely eliminate the positive charge at this residue. We first asked whether the R507H mutant was able to assemble into a ternary complex with PCNA on a CAG extrusion-containing DNA. As judged by pulldown assay (Fig. 5a), at nanomolar protein and DNA concentrations, FAN1 R507H is substantially compromised in complex formation in comparison with the wild type protein. This result suggests a significant weakening of the PCNA-FAN1 binding interface potentially due to loss of H-bond and salt bridge interaction made by Arg507 with Asp232 and Pro253 on PCNA. However, other interactions also likely play a role in stabilizing the FAN1-PCNA complex since it is possible to isolate a FAN1 R507H-PCNA-DNA complex at micromolar concentrations (>75-fold higher concentrations than in the pulldown assay) (see below and Extended Data Fig. 2).

**Figure 5.**
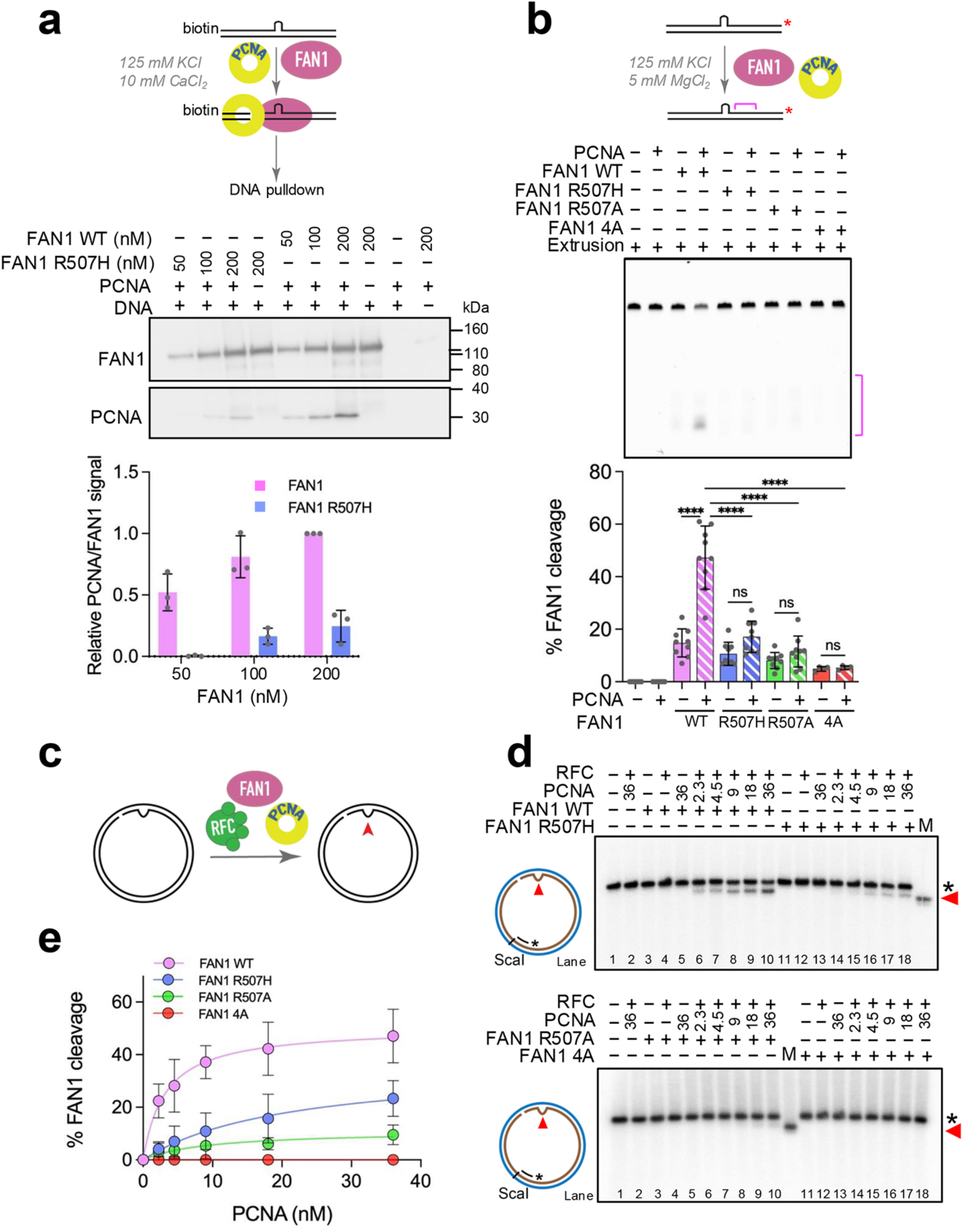
PCNA-dependent FAN1 nuclease activity requires an intact R507 residue. (**a**) Top, schematic of the pulldown assay. Middle, SDS-PAGE analysis of protein complexes bound to magnetic bead-linked DNA substrate harboring a (CAG)_2_ extrusion. Reactions contained PCNA (133 nM as trimer), DNA (50 nM), and indicated concentrations of FAN1 WT or FAN1 R507H. Bottom, Graph represents the ratio of pulled down PCNA relative to FAN1. Data are the average of 3 independent experiments (± SD) and are normalized to the amount of PCNA pulled down with 200 nM FAN1 WT. (**b**) Fifty nM FAN1 WT, FAN1 R507H, FAN1 R507A, or FAN1 4A were incubated with 50 nM of Cy3 labeled DNA substrate harboring a (CAG)_2_ extrusion in the presence or absence of 133 nM PCNA at 125 mM KCl. After 10 min of incubation at 37 °C, reactions were terminated by the addition of formamide to a final concentration of 70 percent. Data are representative of at least three independent experiments (± SD). ***P < 0.001, ****P < 0.0001, ns-P> 0.05, one-way ANOVA with post hoc Tukey’s test. (**c**) Schematic of PCNA-dependent FAN1 nuclease reaction on circular DNA substrate. (**d**) Circular nicked DNA substrate harboring a (CAG)_2_ extrusion (2.5 nM) was incubated in the presence of 10.5 nM RFC, 2.5 nM FAN1 (WT or mutant forms) and PCNA as indicated (see Methods). Reaction products were digested with ScaI, resolved on 1% alkaline agarose gels, followed by hybridization with ^32^P-labeled oligonucleotide probe (Fwd1947). Red arrow indicates the location of the extrusion. (**e**) Quantification of percentage of FAN1 nuclease cleavage from experiments as in (D). Graph based on three independent experiments (± SD). See also Extended Data Fig. 8.

We then evaluated the effect of the R507H substitution on PCNA-dependent FAN1 nuclease activity. At an ionic strength of 125 mM monovalent cation, FAN1 WT as well as the R507H and R507A variants display low levels of nuclease activity (Fig. 5b). Supplementation of the reaction with 133 nM PCNA significantly restored WT FAN1 nuclease activity, in agreement with our previous observations^6^. By contrast, both FAN1 R507H and R507A were refractory to activation by PCNA. Even at much higher PCNA concentrations (up to 1μM), only a modest ∼2-fold stimulation in R507H and R507A nuclease activity was observed (Extended Data Fig. 8a). At low ionic strength (70 mM KCl), the WT FAN1 and FAN1 R507H mutant were indistinguishable in their cleavage efficiency (Extended Data Fig. 8b), suggesting that the R507H substitution does not significantly affect the intrinsic nuclease activity of FAN1. Since Arg507 is located within a conserved region of the protein that harbors a non-canonical PCNA-interacting protein (PIP) box (Extended Data Fig. 8c), we mutated Glu506, Arg507, Ser508, and Val509 residues to Ala. The mutant protein, referred to as FAN1 4A, was refractory to activation by PCNA (Fig. 5b), suggesting compromised ability to form a functional FAN1-PCNA-DNA ternary complex (as observed for the R507H mutant, Fig. 5a).

PCNA can thread onto linear DNA substrates via double-stranded DNA ends (Fig. 1) in agreement with previous studies^35^. However, in cells since such DNA ends are not readily available, PCNA loading requires the catalytic action of the five protein replication factor C complex (RFC)^36,37^. To mimic this physiological situation, we used circular DNA substrates onto which PCNA can be loaded by RFC using a strand discontinuity located 3’ to a (CAG)_2_ extrusion. On these DNA substrates, we observed that, in the absence of DNA-loaded PCNA, FAN1 WT, R507H, R507A, and 4A were all inactive (Fig. 5d, lanes 3-5, 11-13 and Fig. 5e). Increasing concentrations of DNA-loaded PCNA stimulated FAN1 nuclease activity (Fig. 5d, lanes 6-10 and Fig. 5e) for the WT enzyme. By contrast, the PCNA stimulatory effect is strongly reduced for the R507H, R507A, and 4A variants (Fig. 5d, lanes 14-18 and Fig. 5e). The FAN1 activity is restricted to the extrusion-harboring strand (Extended Data Fig. 8d,e). These data suggest that the intact R507 residue plays a critical role in PCNA-dependent FAN1 nuclease activation under physiological conditions, and the R507H variant is severely compromised in its ability to be activated by PCNA on DNAs harboring CAG extrusions. Our findings are consistent with the view that the activation of FAN1 by PCNA, and the efficient removal of CAG extrahelical extrusion by the FAN1-PCNA complex is critical to the ability of FAN1 to stabilize CAG repeat expansion, as has been found in cellular and animal models of HD^10,14^. Our results also predict that HD patients harboring the FAN1 R507H variant are susceptible to higher rates of CAG expansion in the medium spiny neurons of the striatum.

### Structural comparison of FAN1-PCNA-DNA and FAN1 R507H-PCNA-DNA complexes

FAN1 R507H is compromised in its ability to form a ternary complex with PCNA and DNA at molecular concentrations relevant to the nuclease assays discussed above. Therefore, to isolate the FAN1 R507H-PCNA-DNA complex, reactions containing micromolar concentrations of proteins and DNA were assembled as described above for the wild-type protein (Extended Data Fig. 2). The cryo-EM data for this complex also revealed three major 3D classes. The conformations of the mutant ternary complex class 1 and class 2, resembling the two classes of wild-type FAN1 complex, were refined to resolutions of 3.5 Å and 4.0 Å, respectively (Extended Data Fig. 9, 10). After model fitting and refinement, we observed significant alterations in the FAN1 R507H-PCNA interface relative to the wild-type ternary complex. The backbone distance between FAN1 His507 and PCNA Asp232 is 16.5 Å in the mutant complex, while that between FAN1 Arg507 and PCNA Asp232 is 10 Å in the wild-type complex (Fig. 6a,b). Moreover, with His at residue 507, the outer mandible loop of FAN1 undergoes a significant conformational change, resulting in a global shift of FAN1 in respect to PCNA (Fig. 6c,d). The consequence of this displacement is the loss of the stabilizing hydrogen bond and salt bridge with Asp232 and a hydrogen bond with Pro253 on PCNA. Overall, the interface area of PCNA: FAN1 R507H measures 650:748 Å², which is decreased compared to PCNA: FAN1 WT interface (775:816 Å²). In the case of the mutant protein, the interaction is supported by 3 hydrogen bonds contributed by FAN1 T483: PCNA S42, FAN1 H485: PCNA L22, and FAN1 Y514: PCNA E124, as well as 53 van der Waals contacts. These findings suggest that the FAN1 R507H mutation significantly weakens the interaction with PCNA. Additionally, we also observed an allosteric effect of the mutation that results in reconfiguration of the FAN1-DNA interaction (Fig. 6e,f). The DNA complexed with the mutant protein is rotated away from the FAN1 TPR domain towards the VRR_nuc domain (Fig. 6e,f). This movement likely affects the ability of FAN1 R507H to productively associate with the CAG extrusion. Thus, the weakening of the FAN1-PCNA interaction combined with the changes in the FAN1-DNA interface likely contribute to the reduced nuclease activity of FAN1 R507H in the presence of PCNA.

**Figure 6.**
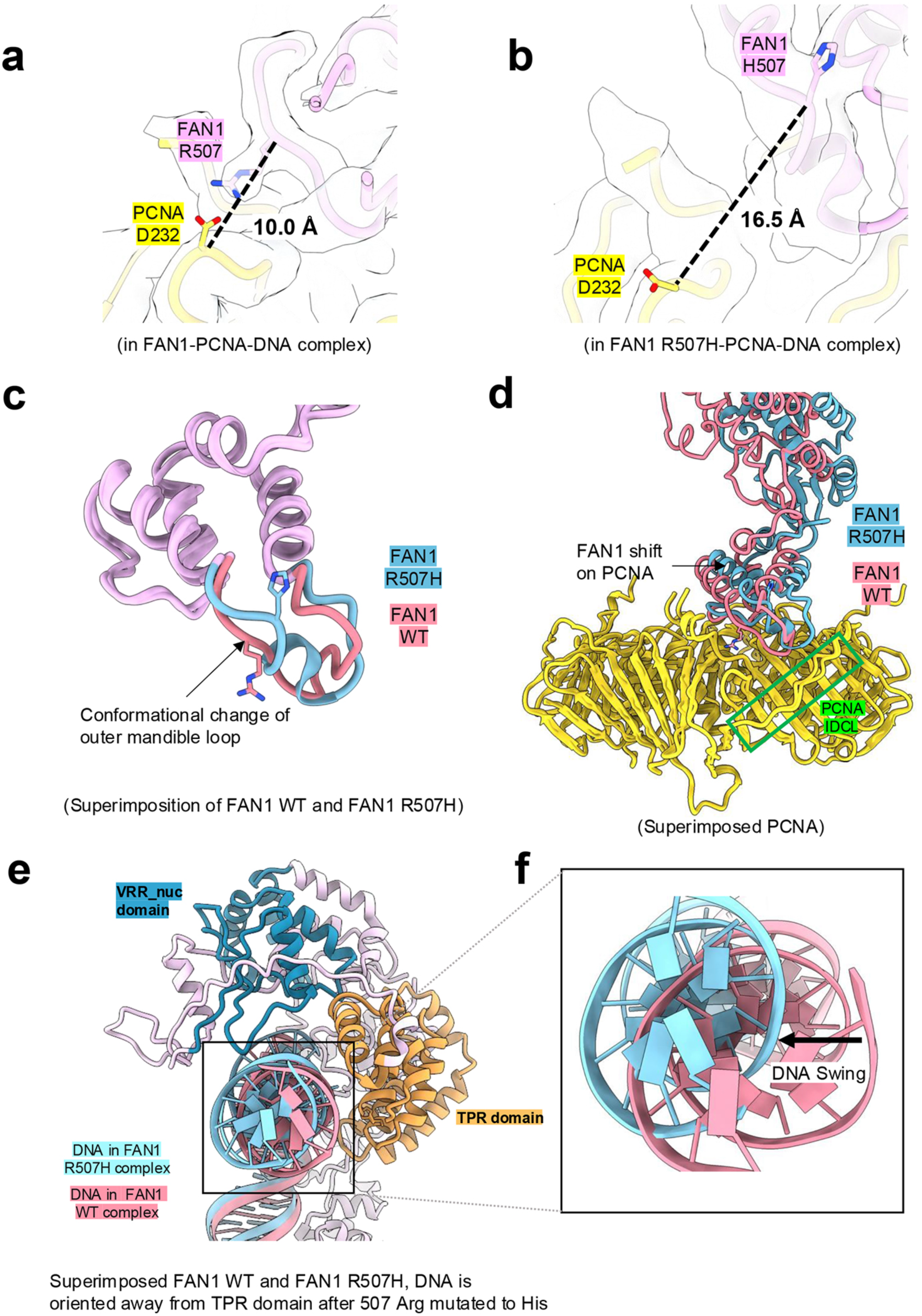
Structural comparison between FAN1-PCNA-DNA and FAN1 R507H-PCNA-DNA. (**a**) Density map overlaid with atomic model of the FAN1-PCNA interface indicating backbone distance of 10.0 Å between FAN1 R507 and PCNA D232. (**b**) Density map overlaid with atomic model of the FAN1 R507H-PCNA interface showing displacement of the polypeptide backbone and the resulting distance of 16.5 Å between FAN1 H507 and PCNA D232. (**c**) Superposition of outer mandible loops of FAN1 WT and R507H indicating conformational changes. (**d**) Displacement of FAN1 relative to PCNA upon mutation of Arg507 to His. (**e-f**) DNA helical axial view showing the DNA tilted away from the TPR domain in the R507H complex relative to WT. See also Extended Data Fig. 9-10.

## DISCUSSION

One of the goals of human biology research is to understand how mutations cause disease. Here, we have used structure-function studies to establish the biological function of a disease-modifying mutation. The importance of the Arg507 residue to FAN1 function is exemplified by the observation that the R507H mutation is one of the strongest modifiers of HD onset^9-11^. This mutation has been also associated with karyomegalic interstitial nephritis, an autosomal recessive disease caused by loss of FAN1 function^38^. Given that the Arg507 residue is not located in the catalytic domain of FAN1, the mechanistic basis of these observations has remained obscure. The cryo-EM structure of the FAN1-PCNA-DNA complex, and the biochemical analyses of FAN1 variants presented here have revealed that Arg507 plays an important role in stabilizing the interaction of FAN1 with PCNA on DNA. Mutation of this residue to His results not only in reconfiguration of the interface between the two proteins, but also alters the position of the DNA in the catalytic pocket (Fig. 6). As a consequence, FAN1 R507H is compromised in its ability to form a FAN1-PCNA-DNA complex under physiological nuclease assay conditions. These effects synergize to render the mutant FAN1 nuclease refractory to activation by PCNA (Fig. 5). Since PCNA-dependent FAN1 nuclease activation results in removal of extrahelical extrusions formed within triplet repeat tracts, attenuation of this activity would be predicted to exacerbate triplet repeat expansion in striatal neurons over the lifetime of the patient, and thereby hasten the disease onset.

The cryo-EM structure of the FAN1-PCNA-DNA ternary complex reveals a unique pincer architecture on FAN1 that is used to latch on to PCNA via an inner mandible and an outer mandible that contains the Arg507 residue. The mandible residues engage with two back to back hydrophobic pockets on PCNA, with the Arg507 considerably stabilizing the interaction via a salt bridge and a hydrogen bond with Asp232 and a hydrogen bond with Pro253. Mutation of Arg507 to His results in reconfiguration of the peptide backbone such that the residue 507 is displaced from Asp232 of PCNA by 6.5 Å. The non-canonical nature of the FAN1-PCNA interaction described here poses the question as to whether the PCNA-binding pincer architecture is unique to FAN1. We therefore evaluated other multi-protein complexes containing PCNA in the Protein Data Bank^39^. Our analysis found that PCNA-Lig1 complex^40^ (PDB:8B8T) is mediated by two loops that resemble the outer and inner mandibles of FAN1 (Supplemental Data Fig. 2). Interestingly, in both cases a non-canonical PIP box is embedded within the outer mandible loop. These observations open up the possibility that the PCNA-binding pincer architecture may be more widespread, especially among proteins that do not have an obvious canonical PIP box.

The collection of conformations identified in our cryo-EM studies suggest intermediate states of the FAN1-PCNA-DNA complex (Fig. 7) wherein the FAN1 protein conformationally samples the DNA substrate (*conformational sampling*), and upon recognition of the (CAG)_2_ extrusion (*lesion scanning*), the catalytic pocket becomes well defined and primed for catalysis (*lesion capture*). In our structure, the CAG extrusion confers a 78° kink to the DNA duplex, similar to the 76° angle observed previously in a structure of FAN1 in complex with double flap DNA^7^. Since extrahelical extrusions produce kinks in the B-form DNA helix^31-33^, we postulate that natural bends in extrusion-harboring DNAs are key conformational features recognized by FAN1. In this model, FAN1 preferentially associates with the kinked DNA conformer and catalyzes its cleavage. Indeed, our cryo-EM studies of the PCNA-FAN1-homoduplex DNA complex display a marked absence of kinked duplex conformation (Fig. 7b and Extended Data Fig. 7); in this case the linear nature of the homoduplex leaves FAN1 with no other choice than to cap the DNA end. That this state is not conducive to nuclease cleavage is supported by the inability of FAN1 to cleave homoduplex DNA, as we have previously shown^6^. Our findings also imply that the degree of the DNA bend at the site of the extrusion dictates the efficiency of FAN1 catalysis, a possibility that we are currently pursuing. It should be noted that end-binding of FAN1 can also be observed on extrusion containing DNAs (Supplemental Data Fig. 3), although these species are not well resolved due to their innate flexibility.

**Figure 7.**
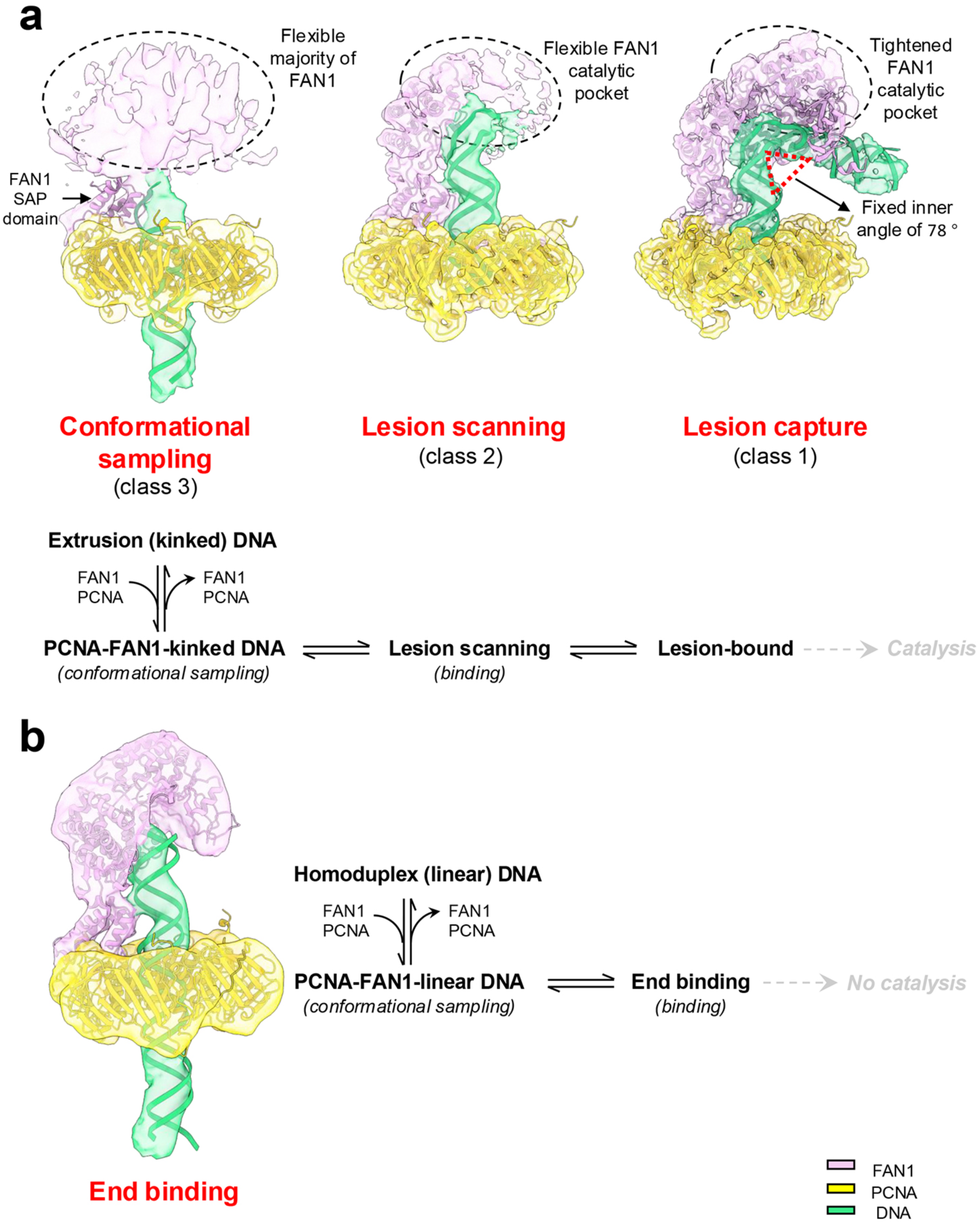
Designation of conformational states of the FAN1-PCNA-DNA complex to steps of FAN1 catalytic cycle. (**a**) Top, comparison of density maps of three classes of the FAN1-PCNA complex bound to DNA substrate harboring a (CAG)_2_ extrusion shown superimposed on atomic models. **Conformational sampling:** In this state, PCNA is well defined and the proximal portion of the DNA is visibly threaded through the PCNA ring. FAN1 and the distal portion of the DNA are poorly defined due to their flexibility. **Lesion scanning**: In this conformation, the proximal end of the DNA is flush with the PCNA ring and is bound by FAN1 (aa 374-609), whereas the distal segment of the DNA from the extrusion site to the 3’ end remains flexible. **Lesion capture:** This state displays PCNA and FAN1 in well-defined form associated with the DNA bent at the extrusion site by an inner angle of 78 °. Bottom, proposed schematic of the FAN1 reaction cycle. FAN1 first samples the DNA substrate for a pre-existing bend such as those found in extrusion-containing DNAs. Upon binding, the FAN1 catalytic pocket undergoes tightening to enable FAN1 to recognize the extrusion site. This primes the DNA extrusion site for catalysis. (**b**) Left, superimposed density map and atomic model of the FAN1-PCNA complex bound to a homoduplex DNA. This conformational state shows a linear DNA, the proximal portion of which is substantially threaded out through the well-defined PCNA ring. The FAN1 protein (in complex with PCNA) caps the distal end of the DNA. Right, proposed reaction sequence of FAN1 on homoduplex DNA. We postulate that the association of the FAN1-PCNA complex with linear homoduplex DNA is conformationally distinct from extrusion containing bent DNA. End-capping by FAN1 is non-productive and does not result in catalysis. See also Extended Data Fig. 8.

The recognition of such bent DNAs harboring extrahelical extrusions by FAN1 also results in reorganization of the FAN1 structure, particularly in the catalytic domain. In the initial substrate-binding mode, the proximal end of the DNA up to the extrusion site is stabilized by the SAP domain of FAN1, whereas the distal portion of the DNA remains flexible as it engages with a highly plastic catalytic domain (*lesion scanning*). The flexibility of the catalytic pocket can also be influenced by the extrusion itself as the location and/or the sequence of the extrusion can vary depending on the pairing with the complementary strand (Supplemental Data Fig. 4). This “wobbling” of the DNA relative to the FAN1 protein facilitates further tightening of the catalytic pocket so as to properly position the DNA substrate at a fixed inner angle (*lesion capture*), thus primed for catalysis.

Previous studies have suggested that PCNA slides along DNA using a cogwheel diffusion mechanism^27,41^. Our structures are consistent with this model and show the DNA threaded through the PCNA ring with a pronounced tilt and with the DNA-protein physical contacts asymmetric with respect to the three PCNA protomers. This mode enables PCNA to interact with the DNA (via sequential hand-off of DNA-protein contacts) while continuously rotating and tilting, thereby resulting in a net forward movement. Taken together with our previous findings that PCNA-activated FAN1 nuclease cleavage is restricted to the DNA strand containing the extrusion^6^, our structures suggest that the cogwheel mode of PCNA translocation along DNA is maintained even with FAN1 nuclease loaded. This enables PCNA-associated FAN1 to track along the helical contour to restrict nuclease cleavage to one DNA strand.

In summary, the structure-function studies described here provide a molecular explanation for (i) the observation that DNA-loaded PCNA is required for activation of the FAN1 nuclease, and (ii) the HD onset hastening effect of the R507H mutation in FAN1. The structure of the FAN1-PCNA complex also lends itself to a model wherein PCNA plays a critical role in activating FAN1-mediated removal of interstrand crosslinks at stalled replication forks. These findings add to the growing view that PCNA functions as an orchestrator of a wide range of DNA repair events in the cell.

## METHODS

### DNA substrate preparation and protein purification

The sequences of the two oligos used for generating the (CAG)_2_ DNA substrate are as follows: Oligo1: 5’-CGA ATT TCT AGA CTC GAG ATC AGC AGC TGC TAG GTC CAG TCT AGA G, Oligo2: 3’-G CTT AAA GAT CTG AGC TCT AGA CGA TCC AGC TCA GAT CTC (see Extended Data Table 1). The annealing protocol followed the method described previously^6^. Recombinant human proliferating cell nuclear antigen (PCNA) was purified from E. coli harboring plasmid pET11a-PCNA as described^6^. Mutants of FAN1 were generated by site directed mutagenesis using the pET28a-His-MBP-FAN1 plasmid as a template. Plasmids for FAN1-R507H (nucleotide change G1520A) and FAN1-Q506A/R507A/S508A/V509A (nucleotide changes C1516G/A1517C/C1519G/G1520C/A1522G/G1523C/T1526C), henceforth referred to as FAN1-4A were synthesized at Genscript. Plasmid for FAN1-R507A mutagenesis on the FAN1-WT plasmid template was carried out with the following oligonucleotide primer pair; FAN1-R507A Fwd: 5’-CAGGTGCAAACGCTAGCTTGCTTCGCCAGTTTCAGAAA-3’,FAN1-R507A Rev: 5’-TTTCTGAAACTGGCGAAGCAAGCTAGCGTTTGCACCTG-3’ with the Agilent QuikChange II Kit according to manufacturer’s instructions. The mutagenesis results in the substitution of C1519 to a G, and G1520 to a C, coding for alanine at amino acid position 507.

FAN1 wild-type (WT), FAN1-R507H, FAN1-R507A and FAN1-4A were expressed and purified from Single step (KRX) cells (Promega Cat# L3002) harboring plasmid pET28a-His-MBP-FAN1 or plasmids expressing the mutant forms of FAN1 as described previously^6^.

### FAN1-PCNA-DNA ternary complex formation

Sixty μL of 15 μM FAN1 wild type-PCNA-DNA combined sample at a ratio of 1:1:1 was incubated at 4°C for 30 min. The combined sample was then subjected to a Superdex 200 Increase 3.2/300 column (GE Healthcare) pre-equilibrated with buffer (25 mM Tris pH 8, 100 mM K-Ac, 0.5 mM TCEP, 10 mM CaCl_2_, 0.2% glycerol) at a flow rate of 60 μL/min. The fractions eluted at 1.1∼1.2 mL were collected for further cryo-EM vitrification (also see Extended Data Fig. 2). The complex of FAN1 R507H-PCNA-DNA and FAN1-PCNA-DNA homoduplex were purified in the same manner as FAN1-PCNA-DNA.

### Cryo-EM sample vitrification and data collection

Two μl of each FAN1-PCNA-DNA complex sample was applied to a 300-mesh copper Quantifoil R 1.2/1.3 + 2 nm holey carbon grid (EMS), which was previously glow-discharged for 90 seconds at 15 mA current using an easiGlow (PELCO). The grids were blotted for 7.5 seconds at a blot force of 2 and immediately vitrified in liquid ethane using a Vitrobot Mark IV (FEI). The clipped grids were screened on a 200 kV Glacios microscope equipped with a Falcon4 detector at Thomas Jefferson University. For high-resolution data collection, grids with FAN1-PCNA-DNA particles were imaged on a Titan Krios microscope operated at 300 kV and equipped with a K3 direct electron detector camera (Gatan) at the National Cryo-Electron Microscopy Facility (NCEF) at the Frederick National Laboratory, MD. A total of 13,068 micrographs of FAN1-PCNA-DNA were collected in super-resolution mode with an energy filter, an image pixel size of 0.537 Å, a nominal magnification of 81,000x, a total dose of 50 e/Å², 40 frames, and a defocus range of −0.75 to −2.5 μm. Additional data collection parameters are provided in Table 1.

The FAN1 R507H-PCNA-DNA complex was vitrified on 300-mesh gold UltrAuFoil R 1.2/1.3 grids, previously glow-discharged for 5 minutes at 20 mA current. These grids were also screened in-house, but data collection was performed at NCEF, using the same K3 detector as for the FAN1-PCNA-DNA dataset. 11,997 movies were collected in counting mode with a pixel size of 0.835 Å, a magnification of 105,000x, a total dose of 50 e/Å², 40 frames, and a defocus range of −0.75 to - 1.75 μm. Additional data collection parameters are provided in Table 1.

The FAN1-PCNA-DNA homoduplex complex was vitrified on 300-mesh gold UltrAuFoil R 1.2/1.3 grids, previously glow-discharged for 5 minutes at 20 mA current. These grids were screened, and 7,809 movies were collected in-house at a magnification of 150,000x with pixel size of 0.95 Å, a total dose of 50 e/Å², 40 frames, and a defocus range of −0.8 to −2.4 μm.

### Cryo-EM single particle analysis of FAN1-PCNA-DNA ternary complex (see Extended Data Fig. 3)

13,068 multi-framed movies of the FAN1-PCNA-DNA complex were imported into CryoSPARC v4.3.1^42^. The output F-crop factor was set to ½ for patch motion correction of the movies, followed by patch CTF (Contrast Transfer Function) estimation with an amplitude contrast of 0.1. Ideal 2D classifications, generated from approximately 1,000 manually picked particles, were used as templates to select particles with a diameter of 150 Å from the entire dataset. After inspection, 13.5 million particles (1,040 particles per micrograph) were accepted into particle pool 1. Forty micrographs, displaying well-distributed particles and good contrast, were selected for particle picking and to train the Topaz picking model. After three rounds of training, the Topaz model, with an average precision of 0.57, was applied for subsequent Topaz particle extraction from the entire dataset. The 3.7 million particles picked by Topaz were combined with particle pool 1. After removing duplicates, 11.9 million particles were extracted with a box size of 256 pixels, Fourier-cropped to 64 pixels. These extracted particles underwent 2D classification, from which 2.6 million particles were selected for generating initial 3D models and further heterogeneous refinement. Four major 3D classes were reconstructed without applying any symmetry. The class with the most complete structure was selected for non-uniform refinement without mask. To continue passing out the ‘garbage particles’, refined particles were expanded by C10, later subjected to ab-initial reconstruction work without symmetry applied to classify 12 classes. Four classes, constituting 40.2% of the total, were selected and around 786 thousand particles remained after removing the duplicates. The remaining particles were re-extracted to 256pix without binning and then non-uniform refined with a solvent mask, yielding a resolution of 3.4 Å with ‘wobbling density’ around the FAN1 catalytic pocket part. To continue analyze the variability, 10 Å resolution was filtered to assess, while simultaneously subjecting the particles to global CTF refinement with Anisotropic Magnification Correction. Then a focused mask was generated to cover the ‘wobbling density’ in order to perform 3D classification with the CTF refined particles under force hard classification. One of the three classes with the most complete structure, containing 262 thousand particles, was selected as the FAN1-PCNA-DNA final state map. The final resolution of 3.8 Å was obtained after a final round of non-uniform refinement with a solvent mask. The cryo-EM map generation of FAN1-PCNA-DNA for the other two classes was similar to the methods above. The local Resolution Estimation with half maps were also performed (see Extended Data Fig. 5).

### Cryo-EM single particle analysis of FAN1 R507H-PCNA-DNA ternary complex (see Extended Data Fig. 9)

11,997 multi-framed movies of the FAN1 R507H-PCNA-DNA complex were imported into CryoSPARC v4.3.1. Patch motion correction was applied for correcting movies without cropping, followed by patch CTF estimation with an amplitude contrast of 0.1. After curating exposures, 10,847 micrographs remained. The 2D templates were generated by importing the volume of the FAN1-PCNA-DNA complex and used as a reference for picking around 6.4 million particles. Topaz picking was also used with the training model (trained after three rounds) to pick about 1.4 million particles from the full dataset. Duplicates were removed after combining the particles picked by templates and Topaz, leaving the remaining particles to be extracted to 256 pixels but binned to 64 pixels. After the first round of 2D classification, 0.77 million particles were selected to reconstruct 3D maps. Four classes were set for ab initio reconstruction, and three classes with a ternary complex appearance were chosen for the next heterogeneous refinement. One of these classes, with the most complete appearance and similar to the FAN1-PCNA-DNA complex, was selected to be extracted to 256 pixels without binning. Non-uniform refinement was carried out to continue refining with the un-binned particles, achieving a resolution of 4.2 Å with blob density in the FAN1 catalytic pocket region. Then, the PCNA ring region was subtracted by creating a localized mask. The subtracted particles then underwent local refinement with another mask localized on the FAN1 catalytic pocket region. The refined model, presenting more continuous density for FAN1, was used as the volume model and input together with the particles before subtraction for non-uniform refinement with a full mask and without particle tilting. The refined particles were further refined by global CTF with Anisotropic Magnification Correction. Finally, the 3D map of the FAN1 R507H-PCNA-DNA final state, with a resolution of 4.0 Å, was generated after a final round of non-uniform refinement with a tight mask. The SPA process to generate the intermediate state followed a similar method to the one described above.

### Cryo-EM single particle analysis of FAN1-PCNA-DNA homoduplex complex (see Extended Data Figure 7)

7,809 multi-framed movies were imported into CryoSPARC v4.3.1. Patch motion correction was applied to correct movies without cropping, followed by patch CTF estimation with an amplitude contrast of 0.1. After curating exposures, 4,733 micrographs remained. 2D templates were generated by importing the volume of the FAN1-PCNA-DNA complex and used as a reference for picking around 1.8 million particles. Topaz picking was also utilized with a trained model (trained after three rounds) to pick approximately 604 thousand particles from the full dataset. Duplicates were removed after combining the particles picked by templates and Topaz, leaving the remaining particles to be extracted to 256 pixels but binned to 64 pixels. After the first round of 2D classification, 1.1 million particles were selected to reconstruct 3D maps. Four classes were set for ab initio reconstruction, and they were subjected to heterogeneous refinement. One of these classes, with a population ratio of 33.4%, was selected for homogeneous refinement without applying a mask. The refined particles were then extracted to 256 pixels but binned to 208 pixels. 2D classification was carried out to further clean the particle pool, after which 292 thousand particles were selected for another round of ab initio reconstruction, setting two classes. One of these classes, with a relatively complete structure, was chosen for two rounds of non-uniform refinements, first without applying a mask and then with applying a tight mask. Finally, the 3D map of the FAN1-PCNA-DNA homoduplex was generated, with a resolution of 6.2 Å, containing 153 thousand particles.

### Model building and refinement

All atomic models presented in this paper were built using Chimera^43^ and Coot^44^. The model refinement was conducted through several rounds of rigid-body, real-space, and B-factor refinement using phenix.real_space_refinement in Phenix V1.20.1^45^. All final models were validated using PDB Validation online server.

### Structure analysis

All ribbon and surface representations were created using ChimeraX^46^. To analyze binding interfaces, PDBsum^26^ was utilized to identify bonding interactions and interatomic distances. A detailed list of interactions at all interfaces can be found in Extended Data Table 2. RMSD values between superimposed PDB structures were calculated using both SuperPose Version 1.0^47^ (superpose.wishartlab.com) and Matchmaker in ChimeraX. Additionally, the Electrostatic Surface Potential was computed and visualized with surface coloring using ChimeraX.

### Data availability

Atomic coordinates for FAN1:PCNA:DNA complex in intermediate, final state, FAN1 R507H:PCNA complex in intermediate, and final state have been deposited in the Protein Data Bank with accession codes 9CG4, 9CL7, 9CHM, and 9CMA, respectively. The cryo-EM density maps have been deposited in the Electron Microscopy Data Bank with accession codes EMD-45568, EMD-45664, EMD-45590, and EMD-45745.

### Nuclease assay

#### FAN1 activity on circular DNA substrates

Assessment of the nuclease activity of FAN1 and its mutants on circular dsDNA substrates was carried out in accordance with protocols established earlier^8^. Briefly, twenty microliter reaction mixes contained 2.5 nM of 3’(CAG)_2_ along with 10.5 nM RFC, 2.5 nM FAN1 or FAN1 mutants as indicated in 20 mM Tris pH 7.6, 0.05 mg/mL BSA, 1.5 mM ATP, 1 mM glutathione (Millipore Sigma, cat # G4251), 5 mM MgCl_2_,125 mM KCl and 2.5 % glycerol. PCNA concentrations were titrated from 2.3 nM, 4.6 nM, 9.2 nM, 18 nM and 36 nM as indicated. Reactions were incubated for 30 min at 37 °C after which they were terminated by the addition of 2 μL of 1 mg/mL Proteinase K (Sigma Aldrich, cat # 3115887001), 140 mM EDTA, 1 % SDS, 1 mg/mL glycogen (Sigma Aldrich cat # 10901393001). Recovered DNA samples were linearized with ScaI and resolved on alkaline gel electrophoresis. The gels were dried at 56 °C for 2 hours in a BioRad Model 583 gel dryer (BioRad Cat # 165-1745), rehydrated and subjected to southern blotting using 5’-^32^P labeled oligonucleotide probes targeting either the extrusion containing strand (Fwd 1947) or the complementary strand (Rev 1975). The gels were exposed to phosphorimager screens and quantified using a Molecular Dynamics PhosphorImager.

#### FAN1 activity on linear DNA substrates

Reactions (20 µL) contained 50 nM of the 3’-Cy3-labeled (CAG)_2_ or homoduplex (or 10nM 3’-radiolabeled +40 nM 3’-Cy3-labeled (CAG)_2_) substrate along with 50 nM FAN1 or FAN1 mutants and 133 nM PCNA (trimer) or as indicated, in 25 mM HEPES KOH pH 7.5, 0.05 mg/mL BSA, 2.5 % glycerol and 5 mM MgCl_2_, and 70 mM KCl or 125 mM KCl. Samples were preincubated with PCNA on ice for five minutes, followed by addition of FAN1 and further incubation at 37 °C for 10 minutes. Reactions were stopped by addition of formamide to a final concentration of 70% and resolved on 15 % urea PAGE gels.

### DNA pulldown assay

Biotinylated DNA substrate harboring a (CAG)_2_ extrahelical extrusion was immobilized on streptavidin beads (Invitrogen, cat # 11205D). Such immobilized DNA was used for the DNA pulldown assays. Briefly, reactions (90 μL) contained 50 nM DNA substrate, 133 nM PCNA (trimer), FAN1 or FAN1 R507H as indicated, 20 mM Tris-HCl pH 7.5, 125 mM KCl, 10 mM CaCl_2_, 0.01% NP40, and 50 μg/mL BSA. First, the DNA was incubated with PCNA on ice for 5 min, followed by addition of FAN1 and further incubation at room temperature for 10 min. Samples were placed on magnet, and the beads were washed 3 times in a buffer composed of 20 mM Tris-HCl pH 7.5, 125 mM KCl, 10 mM CaCl_2_, and 0.01% NP40. Proteins were eluted from the beads by boiling for 5 min at 95 °C in Laemmli buffer and analyzed by SDS-PAGE.

## Acknowledgements

We would like to thank Drs. Gino Cingolani, Ravi R. Iyer, Diane Merry, Richard Pomerantz, Brinda Prasad, Dmitry Temiakov, and Thomas Vogt for helpful discussions. This work was supported by National Institute of Health grants R01 GM144553 and R01 NS118082 and the grant from the Gies Foundation. Initial cryo-EM analysis was carried out at Jefferson Integrated Structural Biology Shared Resources, which is supported in part by grant S10 OD030457. We thank the National Cryo-EM Facility staff at NCI-Frederick for their assistance in data collection. This research was, in part, supported by the National Cancer Institute’s National Cryo-EM Facility at the Frederick National Laboratory for Cancer Research under contract 75N91019D00024.

## Authors contribution

F.L.; A.S.P.; M.B. and A.P. designed experiments. F.L.; A.S.P; M.B. and A.P. performed experiments and analyzed data. C-F. H. was involved in the initial cryo-EM experiments as a manager of the cryo-EM facility at Thomas Jefferson University. R.Y. contributed to cryo-EM data analysis. S.B.; A.M.C.; and S.S., contributed to purification of the mutant forms of FAN1. F.L.; A.S.P.; M.B. and A.P. wrote the manuscript.

## EXTENDED DATA

**Extended Data Fig. 1.**
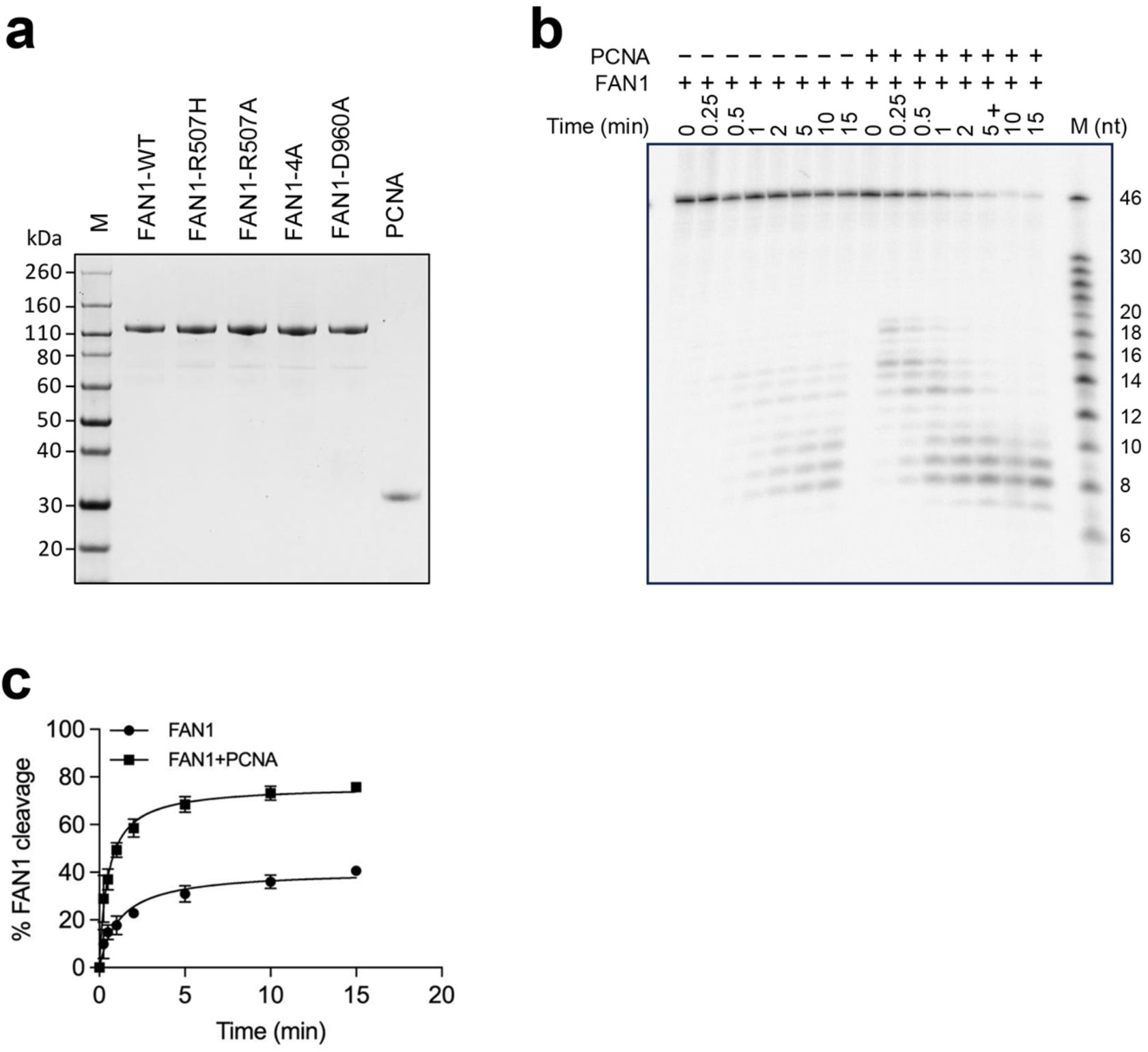
PCNA-dependent FAN1 nuclease activity. (**a**) Electrophoretic analysis of purified proteins used in this study. (**b**) Time course of FAN1 nuclease activity in the presence or absence of PCNA. The part of the gel showing PCNA-dependent FAN1 nuclease activity is shown in Fig. 1a. Here, we show the complete experiment, wherein FAN1 (50 nM) was incubated with DNA substrate harboring a (CAG)_2_ extrusion (50 nM) in the presence or absence of 133 nM PCNA in a buffer containing 125 mM KCl. Samples were withdrawn at indicated time points. (**c**) Quantification of FAN1 nuclease activity based on the data as in (a). Graphs based on 3 independent experiments with error bars representing SD.

**Extended Data Fig. 2.**
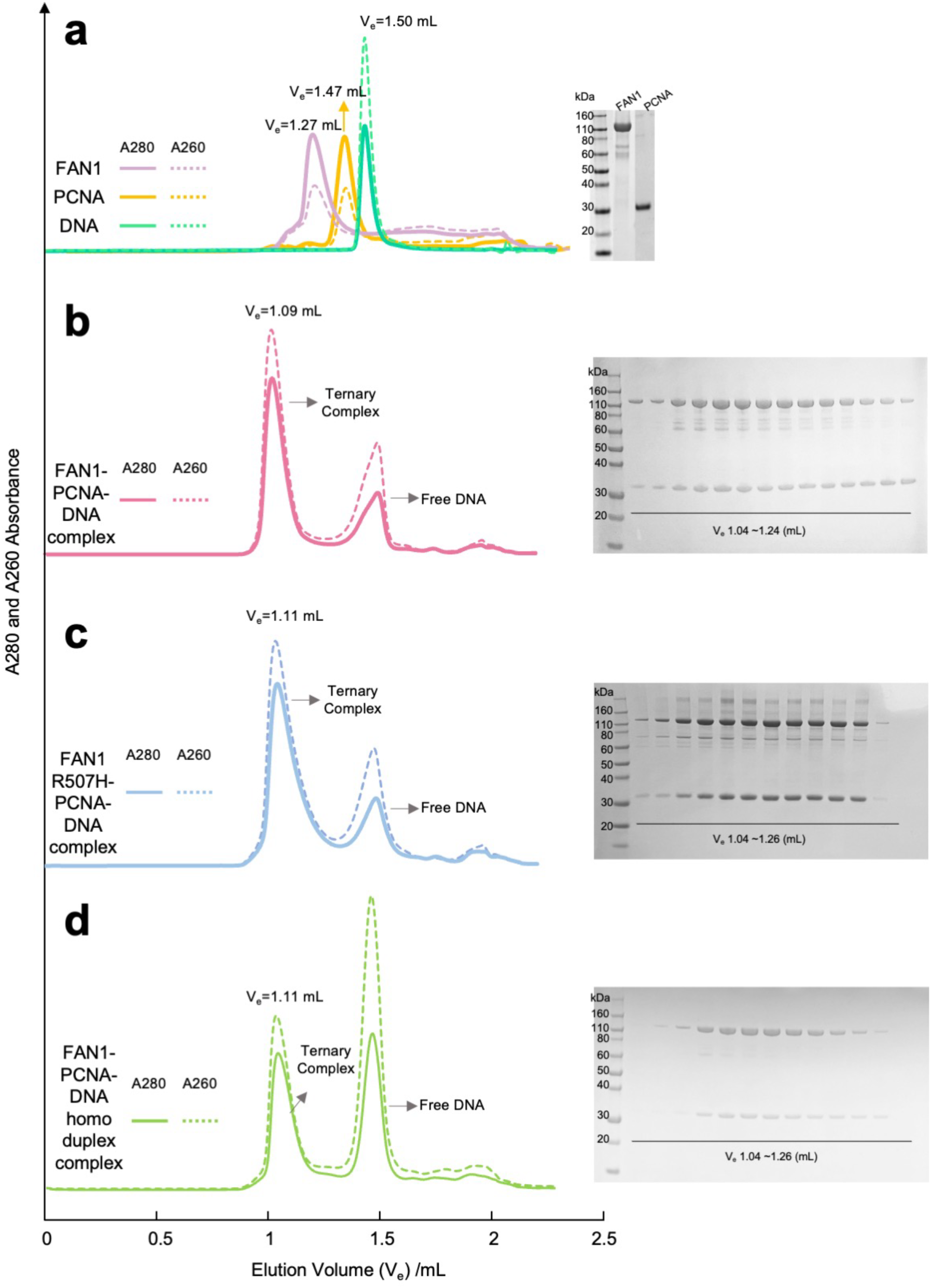
Isolation of FAN1-PCNA-DNA complexes used in this study. Size exclusion chromatographs (left) and SDS-PAGE analysis of eluted fractions are shown for: (**a**) individual proteins or DNA, (**b**) FAN1-PCNA-DNA ((CAG)_2_ extrusion) complex, (**c**) FAN1 R507H-PCNA-DNA ((CAG)_2_ extrusion) complex, (**d**) FAN1-PCNA-DNA (homoduplex) complex.

**Extended Data Fig. 3.**
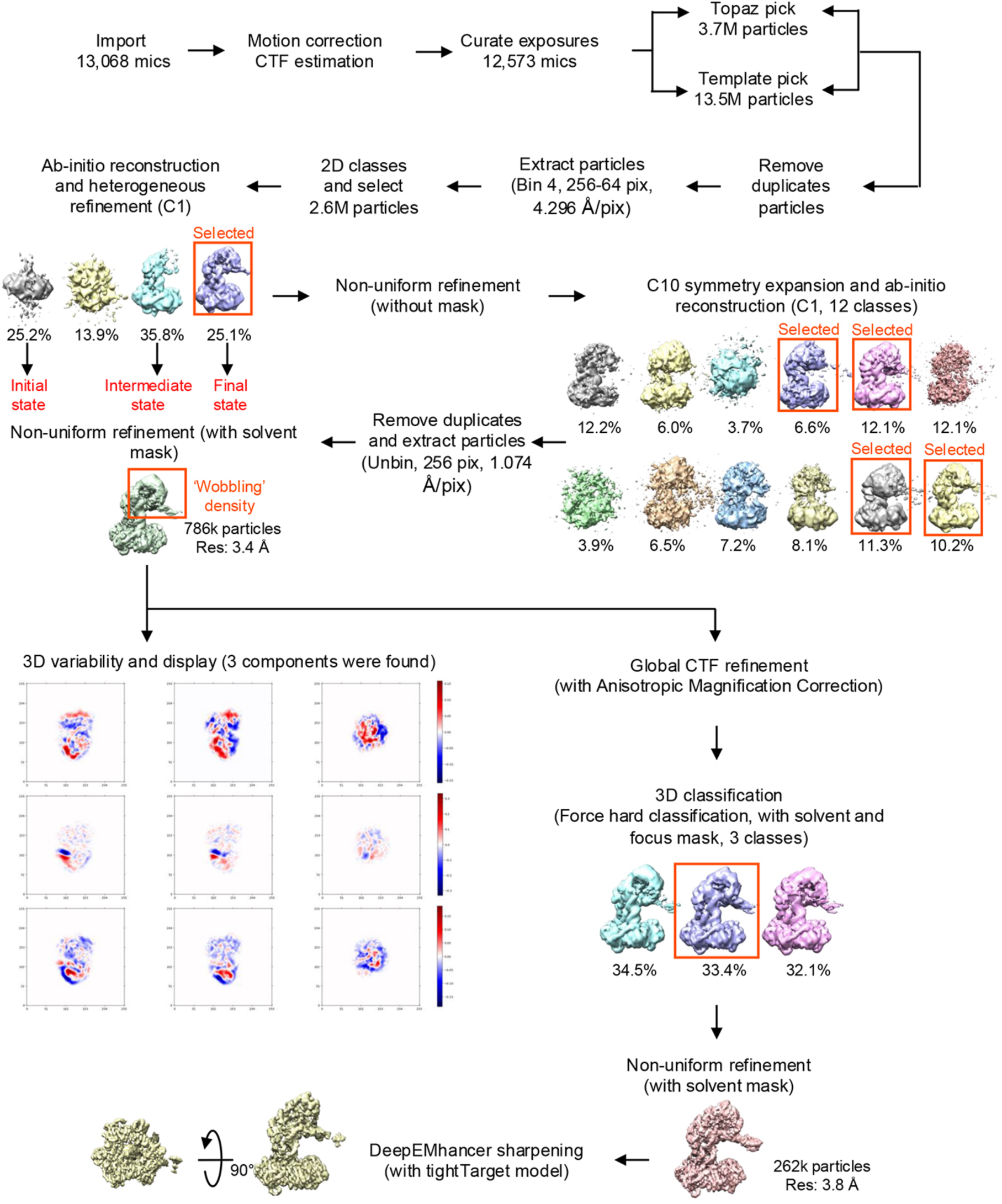
Single Particle Analysis (SPA) processing flowchart for FAN1-PCNA-DNA complex. Also see Methods.

**Extended Data Fig. 4.**
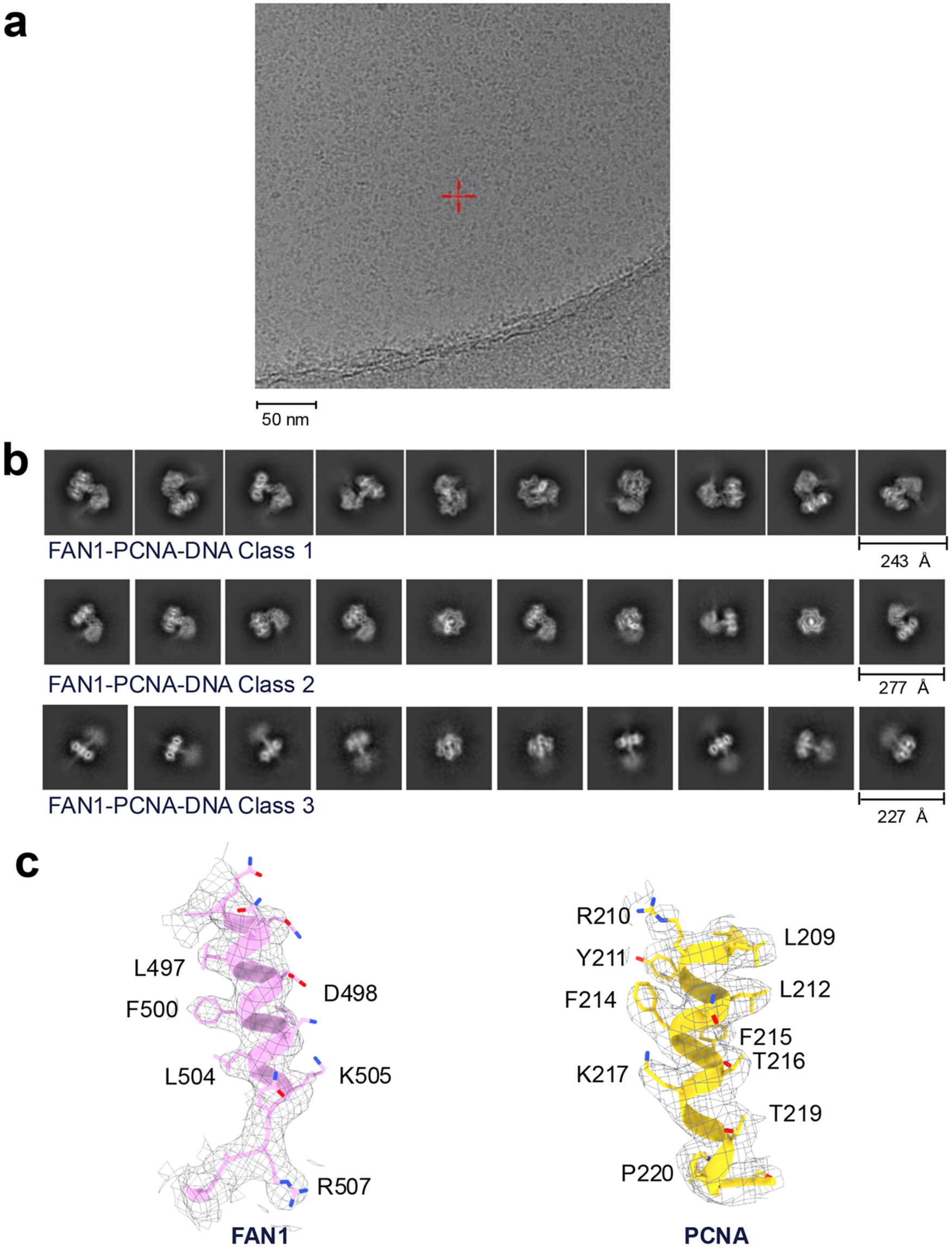
FAN1-PCNA-DNA complex cryo-EM data analysis. (**a**) Representative cryo-EM micrograph of FAN1-PCNA-DNA complex images on a Titan Krios microscope equipped with a K3 detector. (**b**) 2D classifications of three heterogeneous classes. (**c**) Representative regions on FAN1 and PCNA indicating the quality of the 3.83 Å cryo-EM map density.

**Extended Data Fig. 5.**
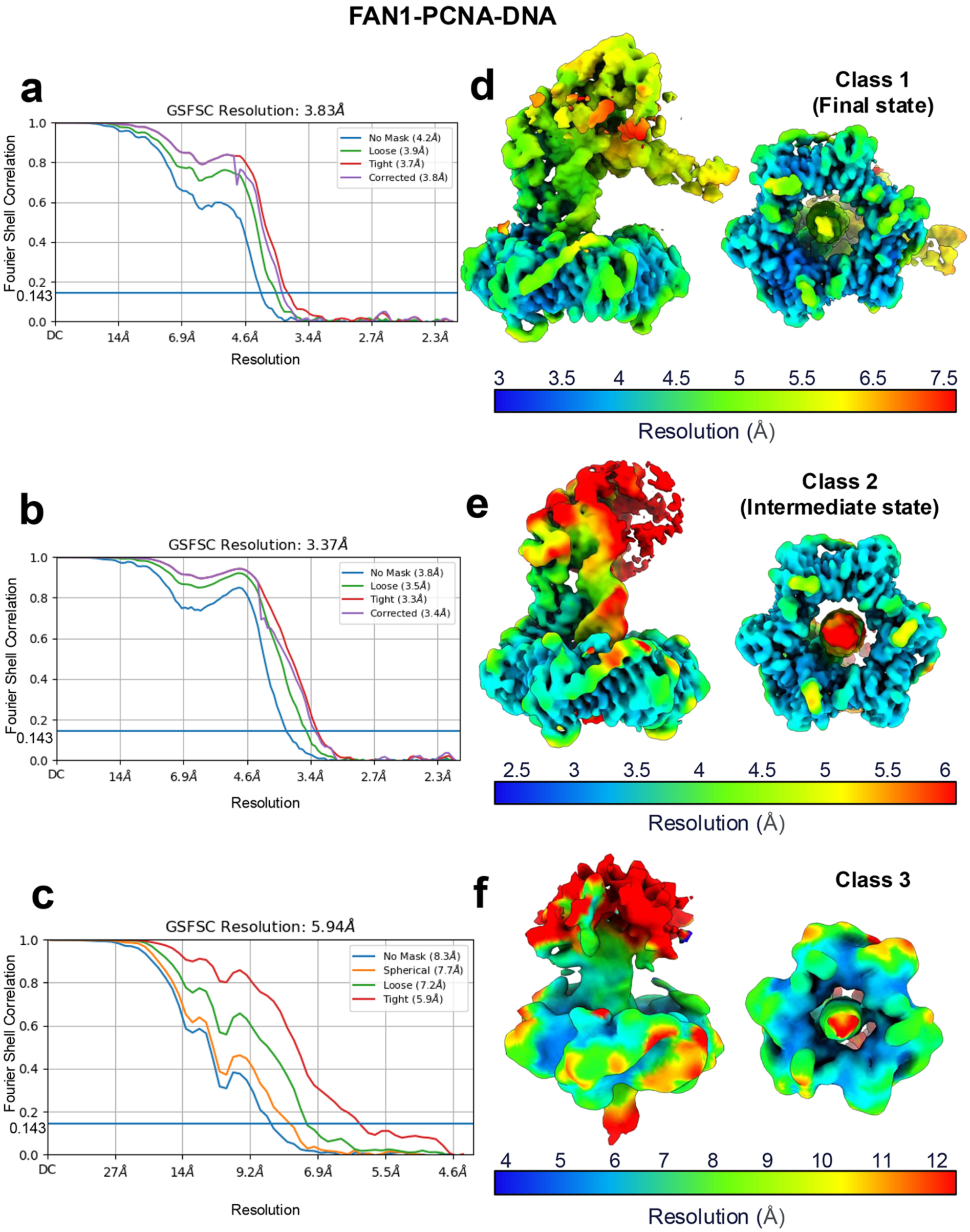
FSCs and localized resolution estimation of FAN1-PCNA-DNA complex. (**a-c**) Fourier Shell Correlation curves with a 0.143 cut-off indicating the estimated resolutions of cryo-EM maps of each of the three classes of the FAN1-PCNA-DNA complex. (**d-f**) Local resolution maps for the three classes.

**Extended Data Fig. 6.**
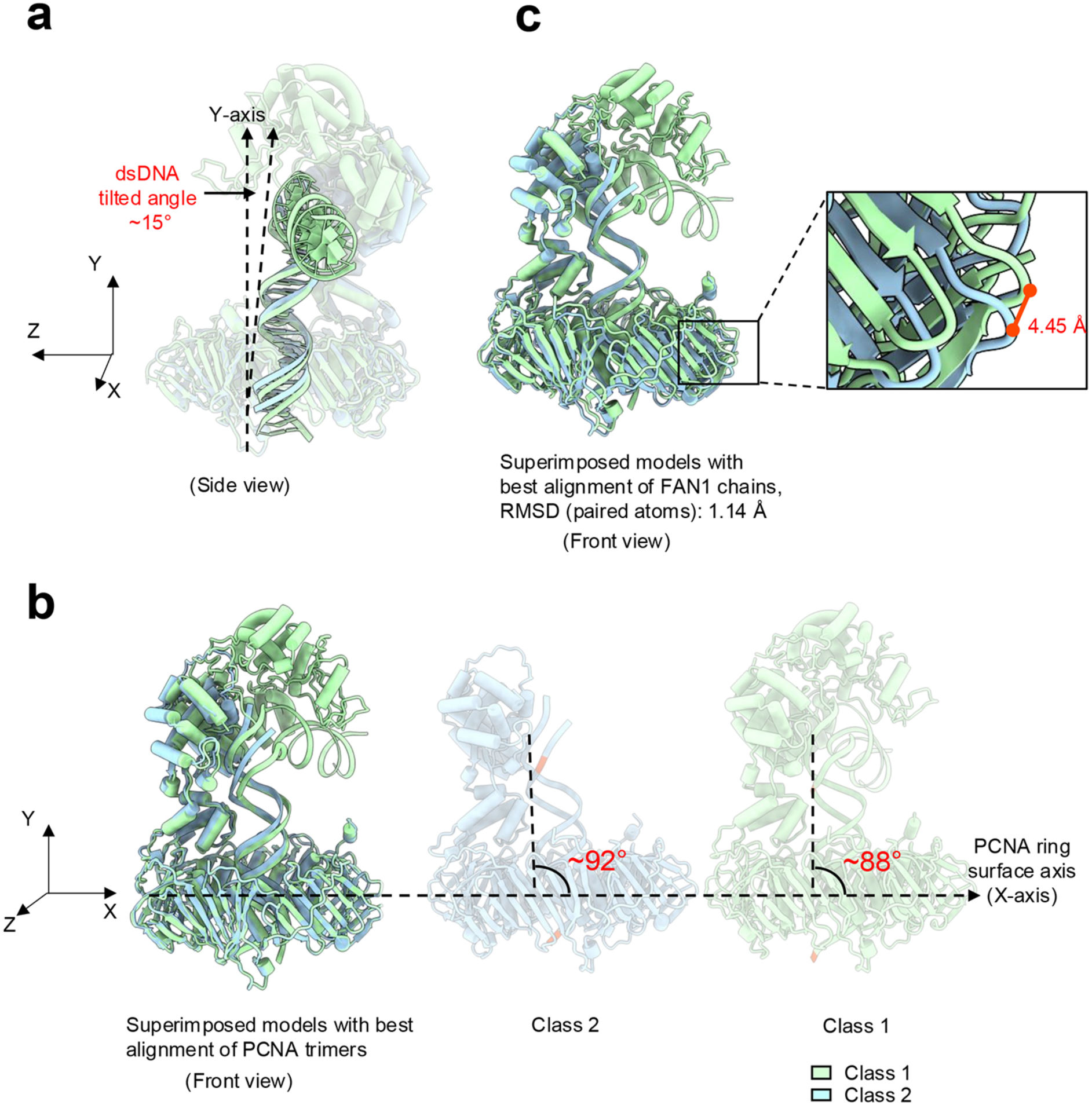
Comparison of conformational differences between class 1 and class 2 states of the FAN1-PCNA-DNA. (**a**) A side-view of the atomic model of the complex shows the dsDNA tilted by ∼15° with regard to Y-axis in an orientation where the PCNA ring sits flat on the X-Z plane. (**b**) Superimposed models of two classes with the best alignment of the PCNA trimers shows the DNA axis tilted by ∼4° in the class 2 conformation relative to class 1. (**c**) Superposition of the two models with best alignment of FAN1 chains. The RMSD between 250 pruned atom pairs is 1.14 Å. The zoomed-in panel on the right shows the region of maximum displacement of PCNA of 4.45 Å.

**Extended Data Fig. 7.**
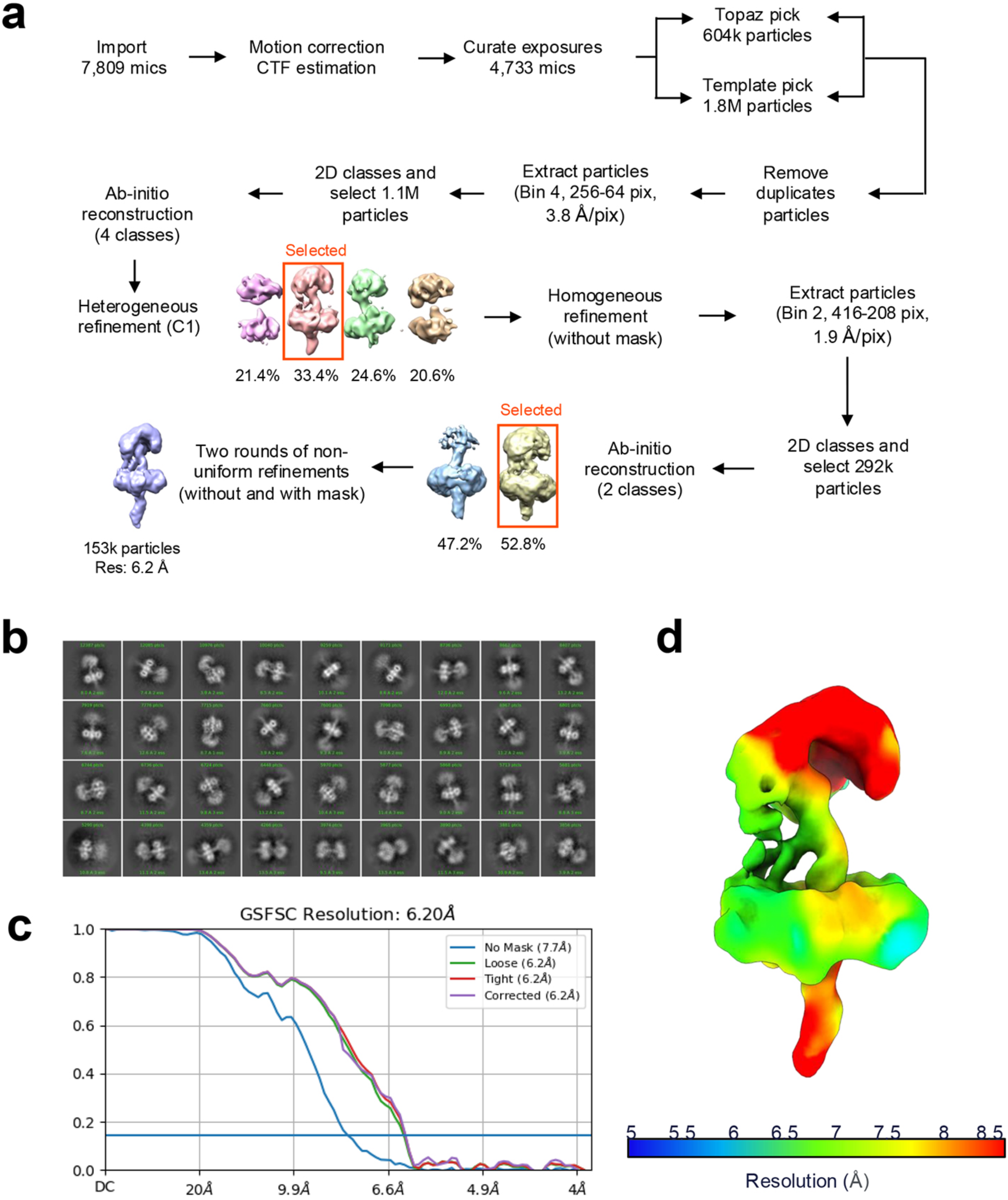
FAN1-PCNA-DNA homoduplex complex cryo-EM data analysis. (**a**) Single Particle Analysis (SPA) processing flowchart for FAN1-PCNA-DNA homoduplex complex. (**b**) 2D classification of FAN1-PCNA-homoduplex DNA complexes. (**c**) FSCs with a 0.143 cut-off indicate the estimated resolution of the cryo-EM maps of FAN1-PCNA-homoduplex DNA. (**d**) Local resolution estimation for the cryo-EM map.

**Extended Data Fig. 8.**
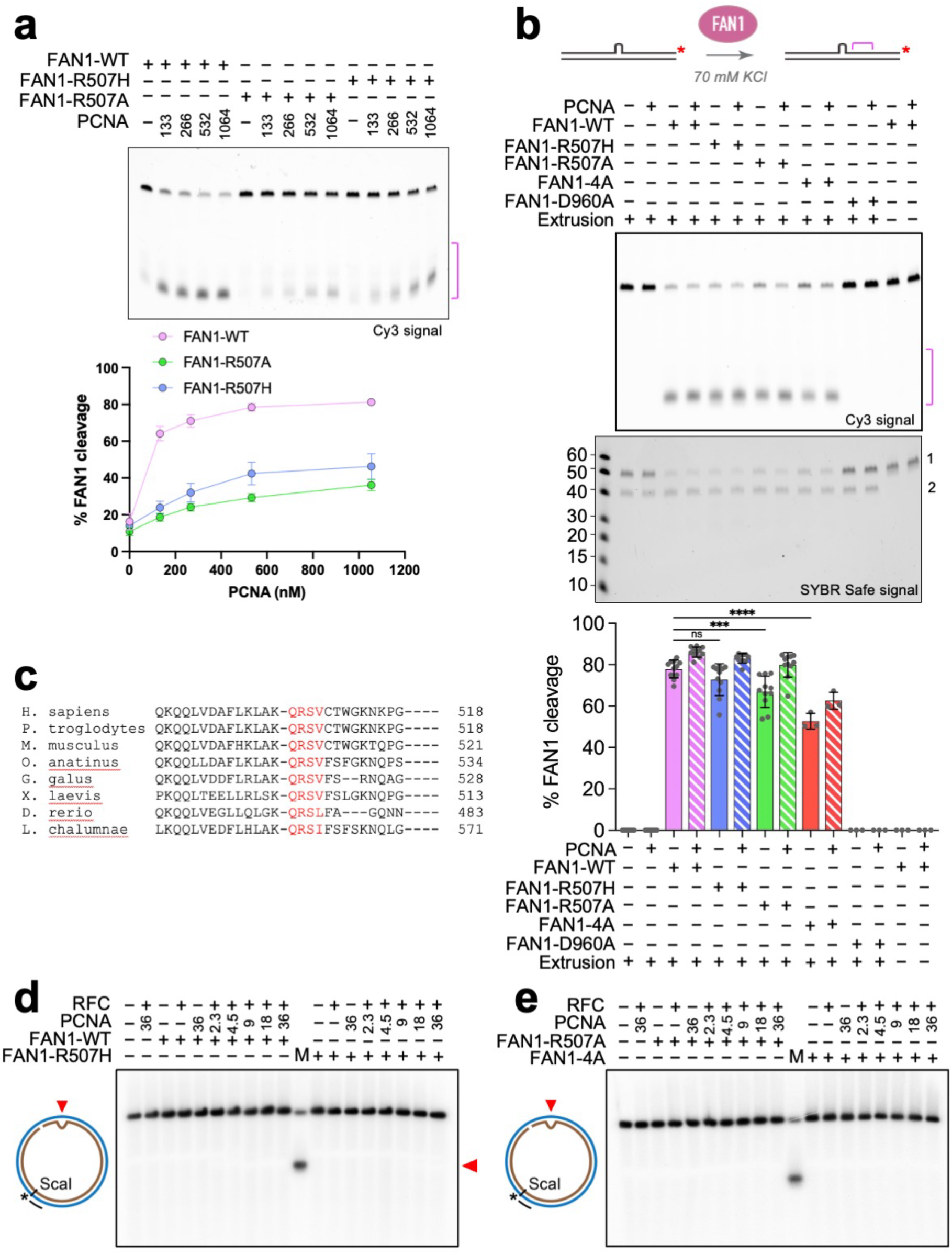
FAN1 mediated cleavage of the 3’(CAG)_2_ substrate. (**a**) Fifty nM FAN1-WT, FAN1-R507H, or FAN1-R507A was incubated with 50 nM of 3’-Cy3 labeled DNA substrate harboring a (CAG)_2_ extrusion in the presence or absence of increasing concentrations of PCNA at 125 mM KCl. After 10 min incubation at 37 °C, reactions were terminated by the addition of formamide to a final concentration of 70 percent. Data are representative of three independent experiments (± SD). The pink bracket indicates FAN1 cleavage products. (**b**) Fifty nM FAN1-WT, FAN1-R507H, FAN1-R507A, or FAN1 4A was incubated with 50 nM of 3’- Cy3 labeled DNA substrate harboring a (CAG)_2_ extrusion in the presence or absence of 133 nM PCNA at 70 mM KCl. After 10 min incubation at 37 °C, reactions were terminated by the addition of formamide to a final concentration of 70 percent and resolved on denaturing polyacrylamide gels (Cy3 signal). The pink bracket indicates FAN1 cleavage products (top). The gel was also stained with SYBR Safe to visualize both DNA strands. (1) DNA 3’-/Cy3/-labeled DNA strand containing (CAG)2 extrusion (cleaved by FAN1 nuclease), (2) complementary strand (not cleaved by FAN1) (middle). Graph represents percentage of FAN1 nuclease activity. Data are representative of at least three independent experiments (± SD). ***P < 0.001, ****P < 0.0001, ns- P> 0.05, one-way ANOVA with post hoc Tukey’s test. (**c**) Sequence alignment of the non-canonical PIP-box (in red) of FAN1 from different species. (**d-e**) 2.5 nM of DNA substrate was incubated in the presence of RFC, PCNA and FAN1 WT or mutant forms as indicated. PCNA concentration was titrated from 2.3 to 36 nM as indicated. Reaction products were digested with ScaI, resolved on 1% alkaline agarose gels, followed by hybridization with ^32^P-labeled oligonucleotide probe (Fwd1975) to visualize the complementary strand. It should be noted that FAN1 displays a cleavage specificity to the extrusion-harboring strand (Fig. 5d), therefore we do not observe FAN1 cleavage of the complementary strand.

**Extended Data Fig. 9.**
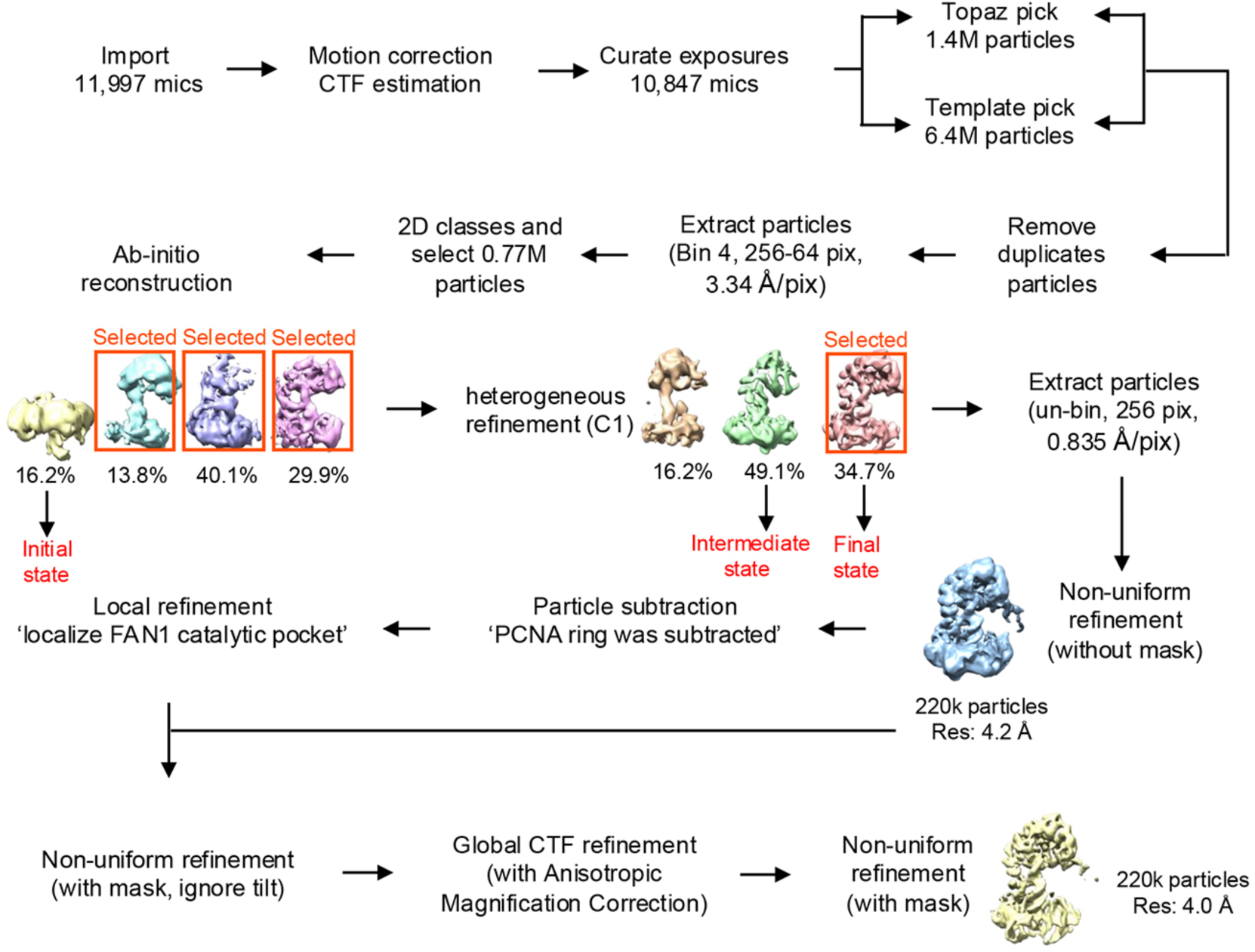
Single Particle Analysis (SPA) processing flowchart for FAN1 R507H-PCNA-DNA complex.

**Extended Data Fig. 10.**
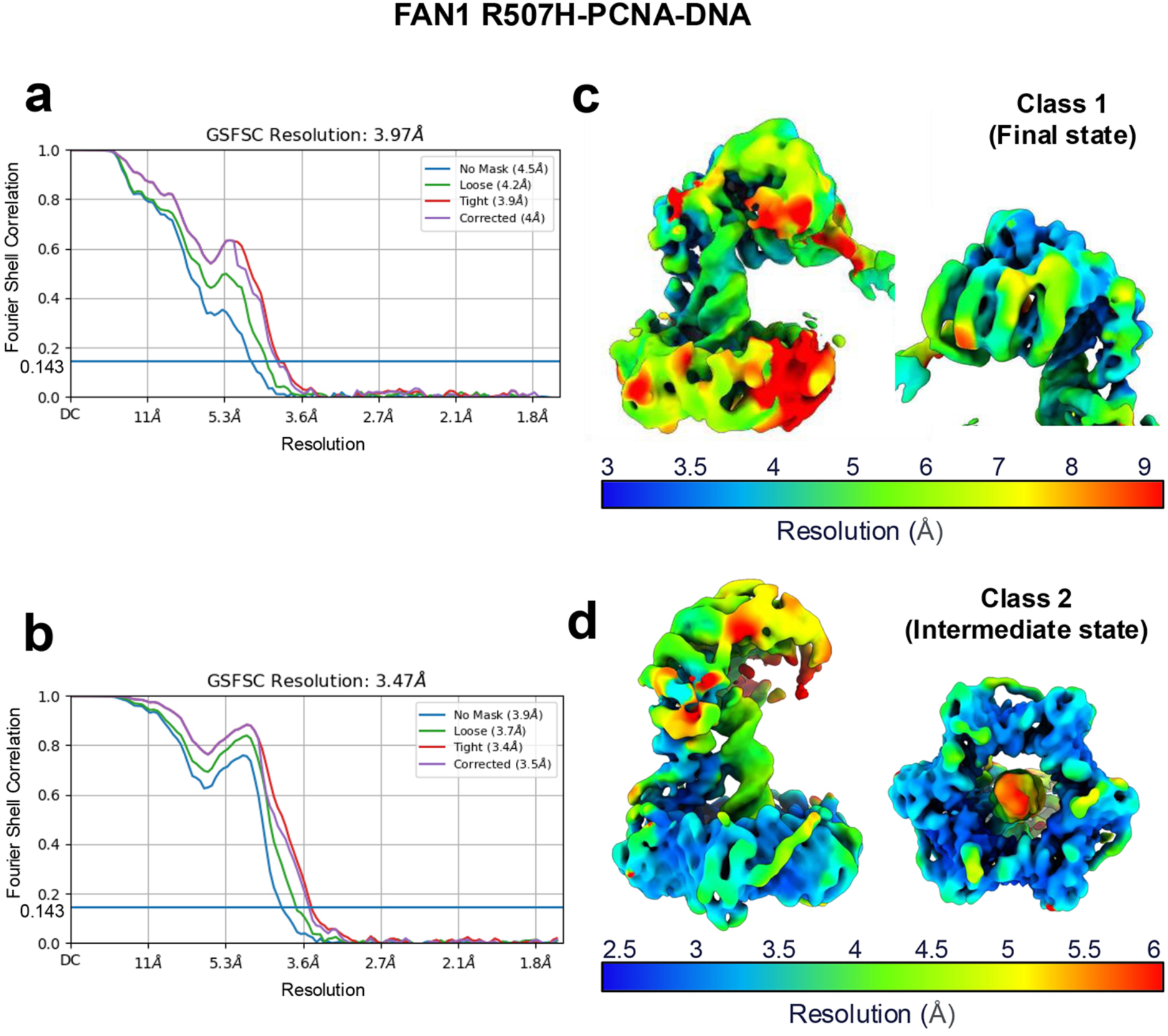
FSCs and local resolution estimation. (**a-b**) FSCs with a 0.143 cut-off show the estimated resolution of the cryo-EM maps of the two FAN1 R507H-PCNA-DNA complex classes. (**c-d**) Local resolution estimation for the two cryo-EM maps.

**Extended Data Table 1.**
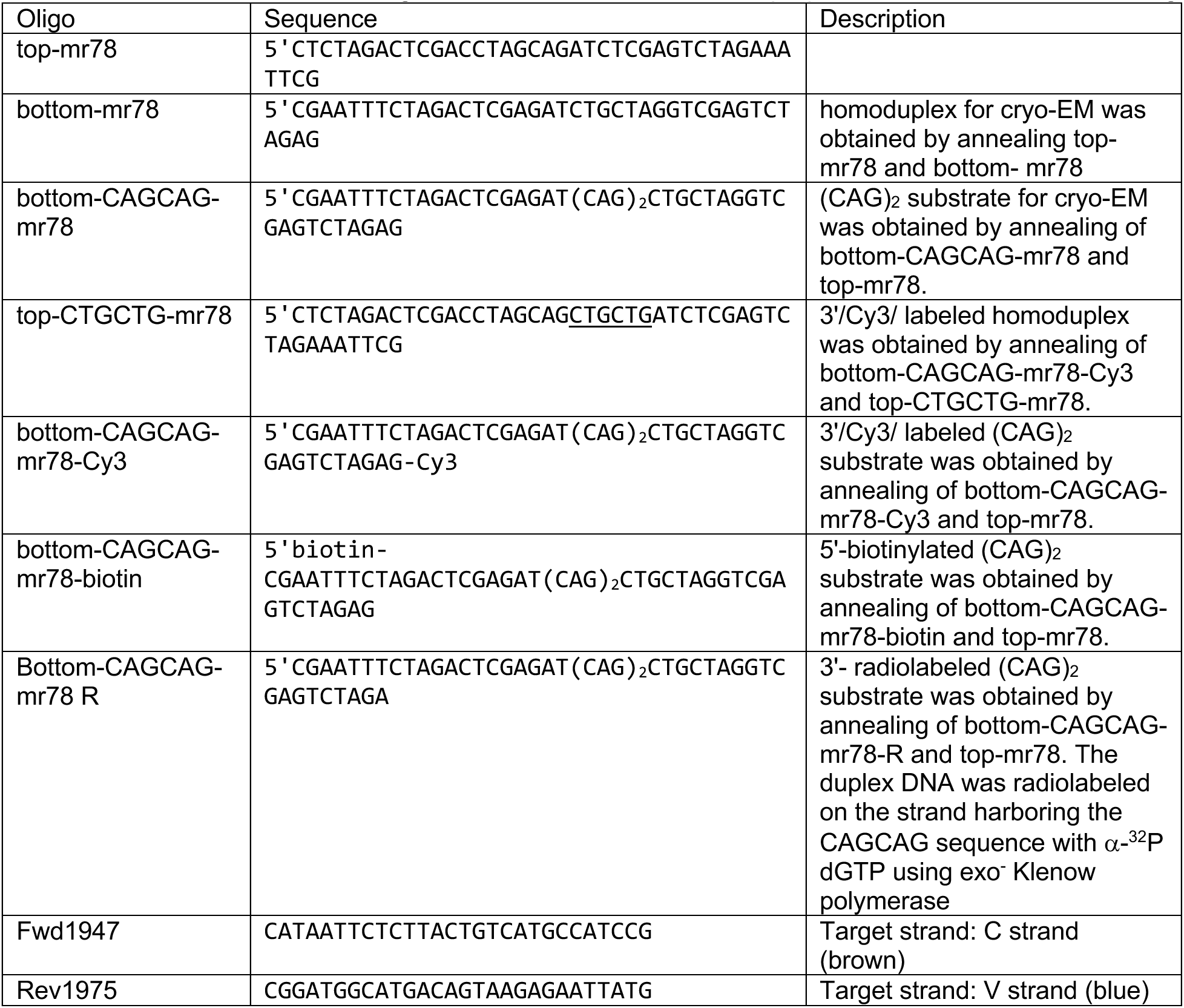
DNA oligonucleotides used for substrate preparation and indirect end labeling.

**Extended Data Table 2.** Atom-atom interactions across FAN1-PCNA interface.

| Interaction Type | PCNA |  | FAN1 |  | Distance (Å) |
| --- | --- | --- | --- | --- | --- |
|  | Atom | Residue | Atom | Residue |  |
| Hydrogen Bonds | OG | 42 SER | O | 483 THR | 2.69 |
|  | OH | 211 TYR | ND1 | 485 HIS | 2.77 |
|  | OD1 | 232 ASP | NH1 | 507 ARG | 2.89 |
|  | O | 253 PRO | NH1 | 507 ARG | 3.21 |
| Salt Bridges | OD1 | 232 ASP | NH1 | 507 ARG | 2.73 |
| Van Der Waals Interactions | OG | 42 SER | C | 483T HR | 3.86 |
|  | CB | 42 SER | O | 483T HR | 3.07 |
|  | OG | 42 SER | O | 483 THR | 2.69 |
|  | OG | 43 SER | CA | 484 PHE | 3.85 |
|  | OG | 43 SER | CB | 484 PHE | 3.68 |
|  | OG | 43 SER | CG | 484 PHE | 3.33 |
|  | OG | 43 SER | CD1 | 484 PHE | 2.81 |
|  | OG | 43 SER | CE1 | 484 PHE | 3.42 |
|  | NH2 | 210 ARG | C | 485 HIS | 3.25 |
|  | NE | 210 ARG | O | 485 HIS | 3.52 |
|  | CZ | 210 ARG | O | 485 HIS | 3.31 |
|  | NH2 | 210 ARG | O | 485 HIS | 2.34 |
|  | CE1 | 211 TYR | O | 485 HIS | 3.87 |
|  | CE1 | 211 TYR | CB | 485 HIS | 3.50 |
|  | CZ | 211 TYR | CB | 485 HIS | 3.34 |
|  | OH | 211 TYR | CB | 485 HIS | 2.68 |
|  | CZ | 211 TYR | CG | 485 HIS | 3.69 |
|  | OH | 211 TYR | CG | 485 HIS | 3.10 |
|  | CE2 | 211 TYR | ND1 | 485 HIS | 3.62 |
|  | CZ | 211 TYR | ND1 | 485 HIS | 3.28 |
|  | OH | 211 TYR | ND1 | 485 HIS | 2.77 |
|  | CA | 22 LEU | CE1 | 485 HIS | 3.81 |
|  | O | 22 LEU | CE1 | 485 HIS | 3.51 |
|  | CD2 | 22 LEU | CE1 | 485 HIS | 3.78 |
|  | O | 21 ASP | NE2 | 485 HIS | 3.47 |
|  | NH2 | 210 ARG | N | 486 LEU | 3.52 |
|  | NH2 | 210 ARG | CA | 486 LEU | 2.98 |
|  | NH2 | 210 ARG | C | 486 LEU | 3.15 |
|  | NE | 210 ARG | O | 486 LEU | 3.51 |
| Van Der Waals Interactions | CZ | 210 ARG | O | 486 LEU | 3.43 |
|  | NH2 | 210 ARG | O | 486 LEU | 2.96 |
|  | OE2 | 258 GLU | CB | 505 LYS | 3.26 |
|  | OE2 | 258 GLU | CG | 505 LYS | 3.24 |
|  | O | 44 HIS | OE1 | 506 GLN | 3.67 |
|  | CB | 44 HIS | OE1 | 506 GLN | 3.84 |
|  | CB | 252 ALA | O | 507 ARG | 3.13 |
|  | O | 253 PRO | CD | 507 ARG | 3.28 |
|  | CG1 | 255 ILE | CD | 507 ARG | 3.37 |
|  | CD1 | 255 ILE | CD | 507 ARG | 3.72 |
|  | CG1 | 255 ILE | NE | 507 ARG | 3.79 |
|  | CD1 | 255 ILE | NE | 507 ARG | 3.53 |
|  | OD1 | 232 ASP | CZ | 507 ARG | 3.22 |
|  | OD1 | 232 ASP | NH1 | 507 ARG | 2.89 |
|  | O | 253 PRO | NH1 | 507 ARG | 3.21 |
|  | CD | 253 PRO | NH1 | 507 ARG | 3.89 |
|  | CG | 232 ASP | NH2 | 507 ARG | 3.21 |
|  | OD1 | 232 ASP | NH2 | 507 ARG | 2.73 |
|  | OD2 | 232 ASP | NH2 | 507 ARG | 3.04 |
|  | O | 44 HIS | CA | 508 SER | 3.80 |
|  | O | 44 HIS | CB | 508 SER | 3.41 |
|  | CB | 44 HIS | CB | 508 SER | 3.89 |
|  | CA | 44 HIS | OG | 508 SER | 3.79 |
|  | C | 44 HIS | OG | 508 SER | 3.45 |
|  | O | 44 HIS | OG | 508 SER | 2.98 |
|  | CB | 44 HIS | OG | 508 SER | 3.04 |
|  | CG | 44 HIS | OG | 508 SER | 3.72 |
|  | ND1 | 44 HIS | OG | 508 SER | 3.54 |
|  | CE | 40 MET | CG2 | 509 VAL | 3.51 |
|  | CB | 47 LEU | CG2 | 509 VAL | 3.68 |
|  | CE | 40 MET | N | 510 CYS | 3.65 |
|  | CE | 40 MET | CA | 510 CYS | 3.84 |
|  | SD | 40 MET | CB | 510 CYS | 3.61 |
|  | CE | 40 MET | CB | 510 CYS | 3.66 |
|  | ND1 | 44 HIS | CB | 510 CYS | 3.22 |
|  | CE1 | 44 HIS | CB | 510 CYS | 3.70 |
|  | SD | 40 MET | SG | 510 CYS | 3.15 |
|  | CE | 40 MET | SG | 510 CYS | 3.78 |
|  | ND1 | 44 HIS | SG | 510 CYS | 3.49 |
|  | CE1 | 44 HIS | SG | 510 CYS | 3.30 |
|  | CG | 234 PRO | CD1 | 512 TRP | 3.79 |
|  | CB | 252 ALA | NE1 | 512 TRP | 3.63 |
|  | CD2 | 126 LEU | CE | 514 LYS | 3.77 |
|  | O | 125 GLN | NZ | 514 LYS | 3.90 |
|  | CD2 | 126 LEU | NZ | 514 LYS | 3.16 |

## References

1. Smogorzewska, A. et al. A genetic screen identifies FAN1, a Fanconi anemia-associated nuclease necessary for DNA interstrand crosslink repair. Mol Cell 39, 36–47 (2010).

2. MacKay, C., et al. Identification of KIAA1018/FAN1, a DNA repair nuclease recruited to DNA damage by monoubiquitinated FANCD2. Cell 142, 65–76 (2010).

3. Yoshikiyo, K., et al. KIAA1018/FAN1 nuclease protects cells against genomic instability induced by interstrand cross-linking agents. Proc Natl Acad Sci U S A 107, 21553–7 (2010).

4. Kratz, K. et al. Deficiency of FANCD2-associated nuclease KIAA1018/FAN1 sensitizes cells to interstrand crosslinking agents. Cell 142, 77–88 (2010).

5. Liu, T., Ghosal, G., Yuan, J., Chen, J. & Huang, J. FAN1 acts with FANCI-FANCD2 to promote DNA interstrand cross-link repair. Science 329, 693–6 (2010).

6. Phadte, A.S. et al. FAN1 removes triplet repeat extrusions via a PCNA- and RFC-dependent mechanism. Proc Natl Acad Sci U S A 120, e2302103120 (2023).

7. Wang, R., et al. DNA repair. Mechanism of DNA interstrand cross-link processing by repair nuclease FAN1. Science 346, 1127–30 (2014).

8. Deshmukh, A.L. et al. FAN1 exo-not endo-nuclease pausing on disease-associated slipped-DNA repeats: A mechanism of repeat instability. Cell Rep 37, 110078 (2021).

9. Consortium, G.M.o.H.s.D. Identification of Genetic Factors that Modify Clinical Onset of Huntington’s Disease. Cell 162, 516–26 (2015).

10. Goold, R. et al. FAN1 modifies Huntington’s disease progression by stabilizing the expanded HTT CAG repeat. Hum Mol Genet 28, 650–661 (2019).

11. Kim, K.H. et al. Genetic and Functional Analyses Point to FAN1 as the Source of Multiple Huntington Disease Modifier Effects. Am J Hum Genet 107, 96–110 (2020).

12. A novel gene containing a trinucleotide repeat that is expanded and unstable on Huntington’s disease chromosomes. The Huntington’s Disease Collaborative Research Group. Cell 72, 971-83 (1993).

13. Zhao, X.N. & Usdin, K. FAN1 protects against repeat expansions in a Fragile X mouse model. DNA Repair (Amst) 69, 1–5 (2018).

14. Loupe, J.M. et al. Promotion of somatic CAG repeat expansion by Fan1 knock-out in Huntington’s disease knock-in mice is blocked by Mlh1 knock-out. Hum Mol Genet (2020).

15. Goold, R. et al. FAN1 controls mismatch repair complex assembly via MLH1 retention to stabilize CAG repeat expansion in Huntington’s disease. Cell Rep 36, 109649 (2021).

16. McAllister, B. et al. Exome sequencing of individuals with Huntington’s disease implicates FAN1 nuclease activity in slowing CAG expansion and disease onset. Nat Neurosci 25, 446–457 (2022).

17. Zhao, X., Lu, H. & Usdin, K. FAN1’s protection against CGG repeat expansion requires its nuclease activity and is FANCD2-independent. Nucleic Acids Res 49, 11643–11652 (2021).

18. Handsaker, R.E. et al. Long somatic DNA-repeat expansion drives neurodegeneration in Huntington disease. bioRxiv, 2024.05.17.592722 (2024).

19. Deshmukh, A.L. et al. FAN1, a DNA Repair Nuclease, as a Modifier of Repeat Expansion Disorders. J Huntingtons Dis 10, 95–122 (2021).

20. Jin, H. & Cho, Y. Structural and functional relationships of FAN1. DNA Repair (Amst) 56, 135–143 (2017).

21. Zhou, W. et al. FAN1 mutations cause karyomegalic interstitial nephritis, linking chronic kidney failure to defective DNA damage repair. Nat Genet 44, 910–5 (2012).

22. Ionita-Laza, I. et al. Scan statistic-based analysis of exome sequencing data identifies FAN1 at 15q13.3 as a susceptibility gene for schizophrenia and autism. Proc Natl Acad Sci U S A 111, 343–8 (2014).

23. Sanchez-Garcia, R. et al. DeepEMhancer: a deep learning solution for cryo-EM volume post-processing. Commun Biol 4, 874 (2021).

24. Abramson, J. et al. Accurate structure prediction of biomolecular interactions with AlphaFold 3. Nature 630, 493–500 (2024).

25. Gulbis, J.M., Kelman, Z., Hurwitz, J., O’Donnell, M. & Kuriyan, J. Structure of the C-terminal region of p21(WAF1/CIP1) complexed with human PCNA. Cell 87, 297–306 (1996).

26. Laskowski, R.A., Jablonska, J., Pravda, L., Varekova, R.S. & Thornton, J.M. PDBsum: Structural summaries of PDB entries. Protein Sci 27, 129–134 (2018).

27. Gonzalez-Magana, A. & Blanco, F.J. Human PCNA Structure, Function and Interactions. Biomolecules 10(2020).

28. De March, M. et al. Structural basis of human PCNA sliding on DNA. Nat Commun 8, 13935 (2017).

29. Georgescu, R.E. et al. Structure of a sliding clamp on DNA. Cell 132, 43–54 (2008).

30. Kochaniak, A.B. et al. Proliferating cell nuclear antigen uses two distinct modes to move along DNA. J Biol Chem 284, 17700–10 (2009).

31. Hsieh, C.H. & Griffith, J.D. Deletions of bases in one strand of duplex DNA, in contrast to single-base mismatches, produce highly kinked molecules: possible relevance to the folding of single-stranded nucleic acids. Proc Natl Acad Sci U S A 86, 4833–7 (1989).

32. Wozniak, A.K., Schroder, G.F., Grubmuller, H., Seidel, C.A. & Oesterhelt, F. Single-molecule FRET measures bends and kinks in DNA. Proc Natl Acad Sci U S A 105, 18337–42 (2008).

33. Shi, X., Beauchamp, K.A., Harbury, P.B. & Herschlag, D. From a structural average to the conformational ensemble of a DNA bulge. Proc Natl Acad Sci U S A 111, E1473–80 (2014).

34. Sinden, R.R. DNA Structure and Function, (Academic Press, San Diego, 1994).

35. Burgers, P.M. & Yoder, B.L. ATP-independent loading of the proliferating cell nuclear antigen requires DNA ends. J Biol Chem 268, 19923–6 (1993).

36. Yao, N. et al. Clamp loading, unloading and intrinsic stability of the PCNA, beta and gp45 sliding clamps of human, E. coli and T4 replicases. Genes Cells 1, 101–13 (1996).

37. Waga, S. & Stillman, B. The DNA replication fork in eukaryotic cells. Annu Rev Biochem 67, 721–51 (1998).

38. Bastarache, L. et al. Phenotype risk scores identify patients with unrecognized Mendelian disease patterns. Science 359, 1233–1239 (2018).

39. Berman, H.M. et al. The Protein Data Bank. Nucleic Acids Res 28, 235–42 (2000).

40. Blair, K. et al. Mechanism of human Lig1 regulation by PCNA in Okazaki fragment sealing. Nat Commun 13, 7833 (2022).

41. Ivanov, I., Chapados, B.R., McCammon, J.A. & Tainer, J.A. Proliferating cell nuclear antigen loaded onto double-stranded DNA: dynamics, minor groove interactions and functional implications. Nucleic Acids Res 34, 6023–33 (2006).

42. Punjani, A., Rubinstein, J.L., Fleet, D.J. & Brubaker, M.A. cryoSPARC: algorithms for rapid unsupervised cryo-EM structure determination. Nat Methods 14, 290–296 (2017).

43. Pettersen, E.F. et al. UCSF Chimera--a visualization system for exploratory research and analysis. J Comput Chem 25, 1605–12 (2004).

44. Emsley, P. & Cowtan, K. Coot: model-building tools for molecular graphics. Acta Crystallogr D Biol Crystallogr 60, 2126–32 (2004).

45. Adams, P.D. et al. PHENIX: a comprehensive Python-based system for macromolecular structure solution. Acta Crystallogr D Biol Crystallogr 66, 213–21 (2010).

46. Pettersen, E.F. et al. UCSF ChimeraX: Structure visualization for researchers, educators, and developers. Protein Sci 30, 70–82 (2021).

47. Maiti, R., Van Domselaar, G.H., Zhang, H. & Wishart, D.S. SuperPose: a simple server for sophisticated structural superposition. Nucleic Acids Res 32, W590–4 (2004).

