## Supplemental Data for "Structural and molecular basis of FAN1 defects in promoting Huntington’s disease"

**Figure 1. Electrostatic surface potential of the FAN1-PCNA-DNA complex.**

**Figure 2. Structural comparison between FAN1 with DNA Ligase 1 in complex with PCNA.**

**Figure 3. The FAN1-PCNA-DNA-FAN1 complex.**

**Figure 4. Possible extrusion isomers of the DNA substrate used in this study.**

#### Supplemental Data Figure 1

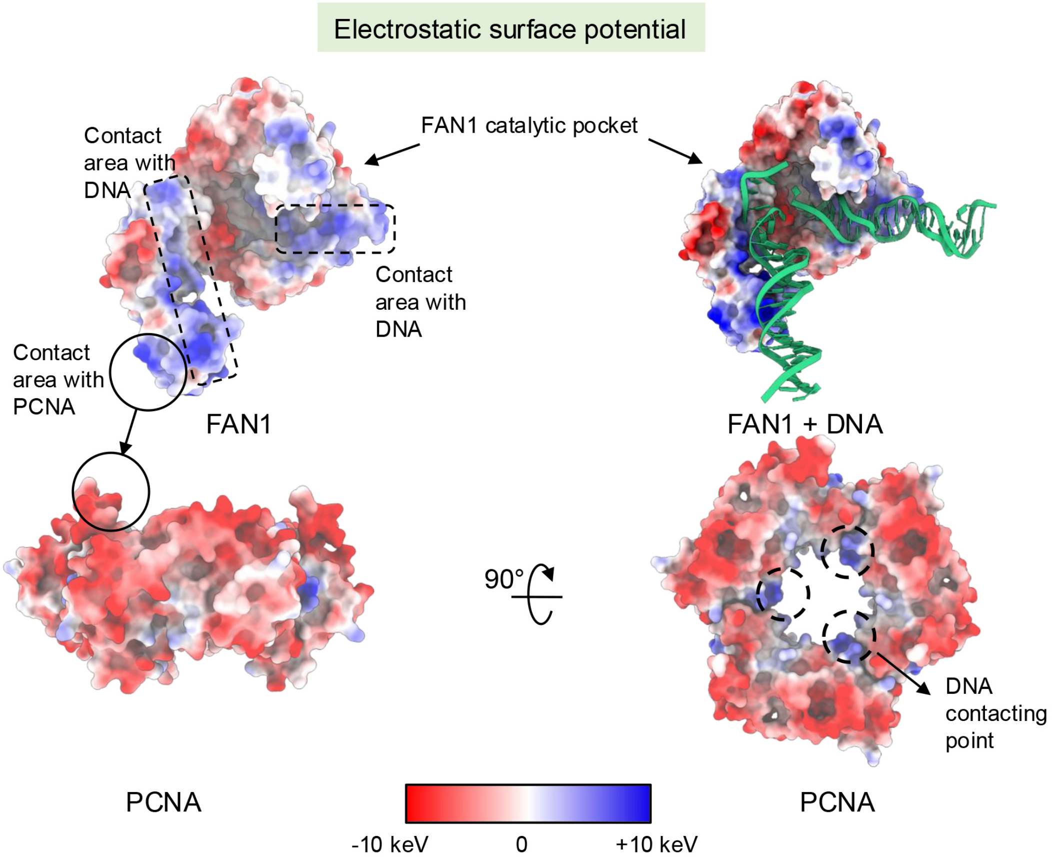


**Supplemental Data Fig. 1. Electrostatic surface potential of the FAN1-PCNA-DNA complex.** The electrostatic surface potential is depicted as a red to blue gradient, with red representing negatively charged regions and blue indicating positively charged regions. The blue regions on FAN1 (top left) are highly positively charged and provide surface suitable for binding negatively charged DNA (top right). Positively charged FAN1 regions (also shown in blue) interact with negatively charged patches on PCNA (shown in red on bottom left). Whereas the outer surface of the PCNA ring is negatively charged (red), the three patches of positive charge are evident on the inner surface of the ring and serve to anchor negatively charged DNA that is threaded through (bottom right).

#### Supplemental Data Figure 2

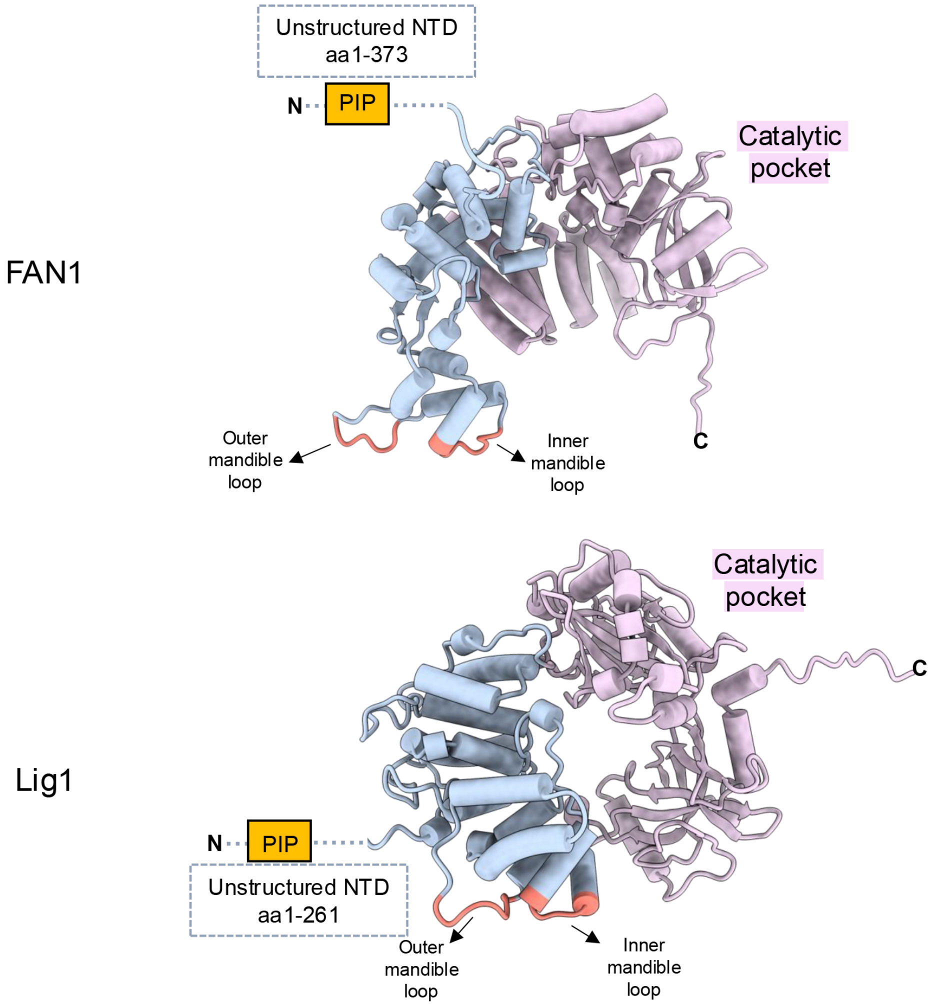


**Supplemental Data Fig. 2. Structural comparison between FAN1 with DNA Ligase 1 in complex with PCNA.**

Atomic models of FAN1 and Lig1 were compared and revealed that LIG1 employs a pincer architecture for PCNA binding that is similar to the pincer architecture at the FAN1-PCNA interface. Lig1 model used here is as per PDB:8B8T.

#### Supplemental Data Figure 3

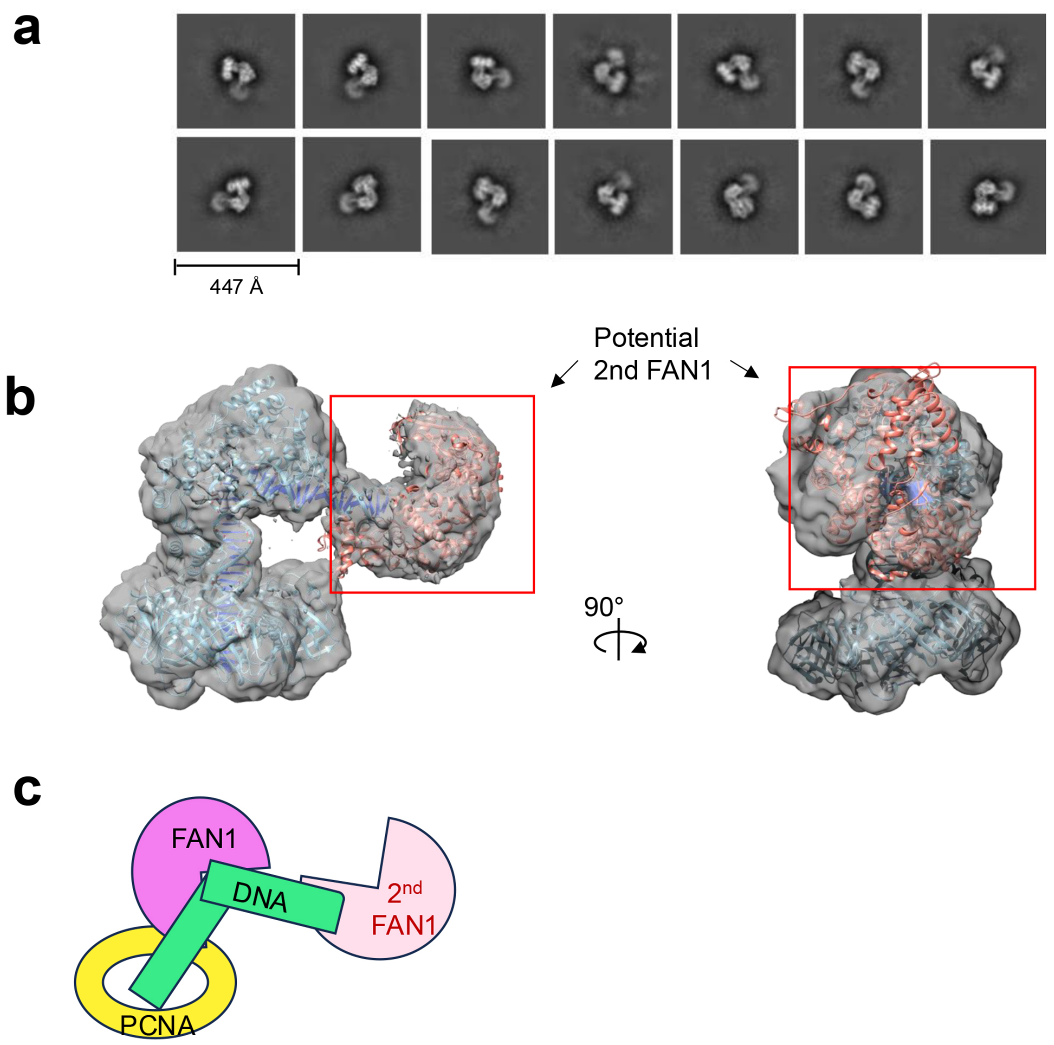


**Supplemental Data Fig. 3. The FAN1-PCNA-DNA-FAN1 complex.**

(**a**) Re-extraction of refined class 1 particles of the FAN1-PCNA-DNA ternary complex (Extended Data Fig. 4) using an enlarged box size (416 pixels, 1.074 Å/pixel) for 2D classification and 3D map generation, revealed an additional flexible density associated with distal end of the DNA substrate. (**b**) The 3D map shows the density corresponding to a putative second molecule of FAN1 (highlighted by the red rectangles) with the atomic models docked within.

(**c**) The cartoon represents a possible assembly arrangement of 2 FAN1 molecules, DNA and PCNA.

### Supplemental Data Figure 4

The switch of nucleotide 'G'

5'-CGAATTTCTAGACTCGAGATCAGCAGCTGCTAGGTCGAGTCTAGAG
3'-GCTTAAAGATCTGAGCTCTA GACGATCCAGCTCAGATCTC

5'-CGAATTTCTAGACTCGAGATCAGCAGCTGCTAGGTCGAGTCTAGAG
3'-GCTTAAAGATCTGAGCTCTAG ACGATCCAGCTCAGATCTC

Extrusion site

Cleavage site

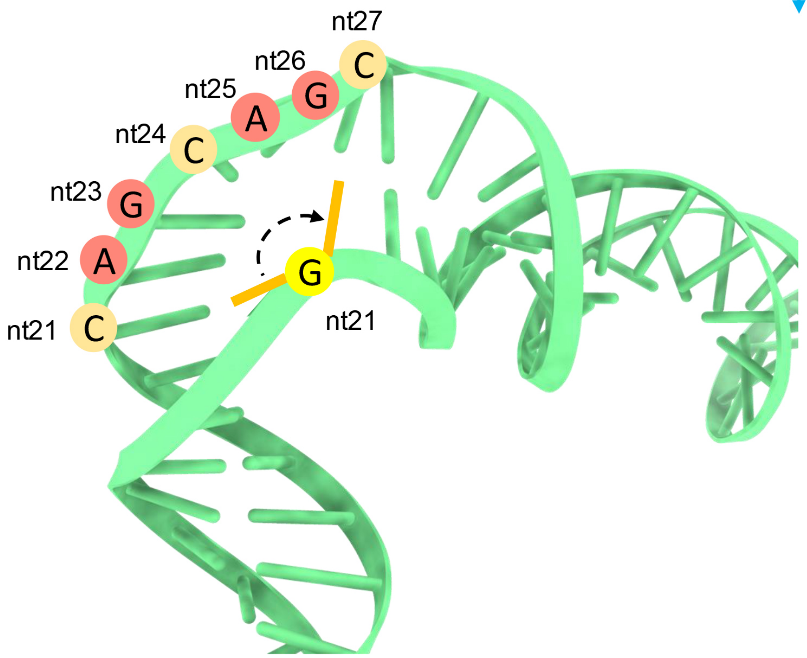


**Supplemental Data Fig. 4. Possible extrusion isomers of the DNA substrate used in this study.**

The G residue on the complementary strand (highlighted in yellow) can pair either with $C_{21}$ or $C_{27}$. Therefore, we cannot rule out the possibility that the complexes described in this study may be composed of two isomers of the DNA substrate that harbor either a $(CAG)_2$ extrusion or an $(AGC)_2$ extrusion.
